# Microbial diversity and community dynamics in an active, high CO_2_ subsurface rift ecosystem

**DOI:** 10.1101/2022.10.24.511890

**Authors:** Daniel Lipus, Zeyu Jia, Megan Sondermann, Robert Bussert, Alexander Bartholomäus, Sizhong Yang, Dirk Wagner, Jens Kallmeyer

## Abstract

The Eger Rift subsurface is characterized by frequent seismic activity and consistently high CO_2_ concentrations, making it a unique deep biosphere ecosystem and a suitable site to study the interactions between volcanism, tectonics, and microbiological activity. Pulses of geogenic H_2_ during earthquakes may provide substrates for methanogenic and chemolithotrophic processes, but very little is currently known about the role of subsurface microorganisms and their cellular processes in this type of environment. To assess the impact of geologic activity on microbial life, we analyzed the geological, geochemical, and microbiological composition of rock and sediment samples from a 240 m deep drill core, running across six lithostratigraphic zones. In addition, we evaluated diversity as well as metabolic attributes of bacterial and archaeal communities. Our investigation revealed a distinct low biomass community, with a surprisingly diverse Archaea population, providing strong support that methanogenic archaea reside in the Eger subsurface. Geochemical analysis revealed sulfate and sodium concentrations as high as 1000 mg L^−1^ in sediment samples from a depth between 50 and 100 m and in weathered rock samples collected below 200 m.

Most microbial signatures could be assigned to common soil and water bacteria, which together with the occurrence of freshwater Cyanobacteria at specific depths, emphasize the heterogenous, groundwater movement driven nature of this terrestrial subsurface environment. Although not as frequently and abundantly as initially expected, our investigations also found evidence for anaerobic, autotrophic, and acidophilic communities in Eger Rift sediments, as sulfur cycling taxa like *Thiohalophilus* and *Desulfosporosinus* were specifically enriched at depths below 100 m. The detection of methanogenic, halophilic, and ammonia oxidizing archaeal populations demonstrate that the unique features of the Eger Rift subsurface environment provide the foundation for diverse types of microbial life, including the microbial utilization of geologically derived CO_2_ and when available H_2_, as a primary energy source.

## Introduction

The deep biosphere represents one of the largest and most unique ecosystems on earth. Subsurface regions belonging to these systems are estimated to comprise up to one third of the total global biomass and harbor a diverse array of geochemical settings and microbial habitats [1, 2]. While microbial life in marine subsurface sediments is relatively well constrained [3, 4], assessments of indigenous microbial communities and their contribution to biochemical cycles in deep terrestrial sediments and hard (i.e. igneous) rocks remain limited. As Earth’s terrestrial subsurface also represents an essential source of groundwater, minerals, metals and hydrocarbons, geochemical and microbial explorations into these systems are of high scientific and industrial relevance. Although the often extreme conditions (no or little oxygen, no light, temperatures of more than 100° C, high pressures, limited carbon sources, and sometimes no water) are believed to push life to its limit and significantly decrease microbial turnover [5], previous studies have suggested Earth’s subsurface to harbor an unexpected phylogenetic diversity and accommodate a variety of unknown microbial populations, often referred to as microbial dark matter [6-9]. Explorations of metabolic processes in subsurface settings have asserted the importance of chemolithoautotrophic, and organotrophic lifestyles as well as methane cycling [10], emphasizing the dependence on geochemically and abiotically derived substrates such as H_2_ and CO_2_. Due to the scarcity of bioavailable carbon, heterotrophic communities in deep continental subsurface settings are believed to rely on the production of fixed carbon through these primary processes.

Subsurface environments characterized by high CO_2_ conditions have become research areas of particular scientific and industrial interest, as both artificial and natural reservoirs are being studied to evaluate the effect of geologic carbon sequestration, capture and storage (CSS), a process in which anthrophonic CO_2_ is injected into subsurface structures for storage [6, 11-13]. Besides influencing geochemical processes in the subsurface, lowering the pH, causing greater weathering and mineral dissolution, elevated levels of CO_2_ can significantly impact the microbial biosphere. This means CO_2_ can interrupt cellular processes, such as protein synthesis and DNA replication, effectively acting as a sterilizing agent. On the contrary, CO_2_, can spark microbial growth by directly acting as a substrate for autotrophic growth or dissolving and providing minerals and nutrients from the surrounding environment. Explorations into microbial distribution, diversity and metabolisms have accentuated the impact of CO_2_ in the terrestrial subsurface, showing microbial populations in such settings to be at least temporarily reduced [11, 12, 14] and enriched in chemolithotrophic iron and sulfur oxidizing bacteria as well as methanogenic archaea. A limited number of metagenomic studies have evaluated the metabolic capabilities of CO_2_ adapted subsurface microbial communities. Freedman et al. [13] was able to recover *Thiobacillus, Gallionella*, and Hydrogenophales draft genomes from 2600 m McElmo Dome aquifer samples, reconstructing metabolic pathways from carbon fixation via the Wood Ljungdahl pathway and Calvin Benson cycle, sulfur oxidation and partial nitrate reduction. Work from Emerson et al. [6] and Probst et al. [15], confirm the relevance of these carbon fixation pathways, as similar genes were found to be enriched in 46 different genomes recovered from CO_2_ saturated Crystal Geyser subsurface fluids. In addition, Gulliver et al. [12], and Trias et al. [16] also reported the metabolic potential for carbon fixation and sulfur oxidation by *Sulfuricurvum* and members of the Comamonadaceae and detected an increased number of methanogenesis genes in CO_2_ injected subsurface aquifers. While analyses to this date provide strong evidence that microbial communities can at least temporarily adapt to high CO_2_ conditions, only few efforts have been made to evaluate microbial life in natural environments with continuous, long term CO_2_ exposure. Thus, additional work covering a wider range of naturally occurring geological and geochemical high CO_2_ subsurface regions, may provide further insights on the direct and indirect impact of CO_2_ on microbial distribution and behavior and long-term development of such ecosystems.

The geodynamically active Eger Rift region in West Bohemia (Czech Republic) is part of the Počatky-Plesná Fault Zone (PPZ) and characterized by a unique combination of CO_2_ rich mantle degassing and regular seismic activity. Frequent earthquake swarms and high flow rates of mineral rich fluids create a distinct sedimentary composition make this region an excellent study site for the evaluation of microbial distribution, abundance, and processes under unusual deep subsurface conditions. Periodically occurring earthquake swarms stimulate the migration of geogenic CO_2_ from active magma chambers at the crust-mantle boundary and from lithospheric mantle depths of about 65 km [17-19] resulting in CO_2_ rich sediments and the formation and accumulation of CO_2_ in groundwater structures such as saline aquifers [20]. At the surface, CO_2_-rich gas is discharged in the form of natural cold gas exhalation systems such as mofette sites or released at mineral water springs [21-26]. The exceptional geo- and physicochemical conditions are likely to affect microbial development and activity, may foster microbial processes through elevated substrate support [27], and potentially trigger a diverse range of interesting rock-fluid interactions as part of geodynamic processes in the lithosphere [18, 26, 28]. On the contrary, elevated CO_2_ levels can cause hypoxia and acidification of the soil, the mobilization of metals, and may drive life to its limit, leaving the ability of microbial communities to survive in this subsurface environment up for debate.

Several efforts have been made to study microbial life and the microbial responses to geological processes in the Eger Rift region. A 2005 study by Bräuer et al [19] described the discovery of elevated concentrations of H_2_ and biogenic methane in mineral spring waters following a seismic event, suggesting that abiotic derived substrates can provide the foundation for microbial life in the Eger Rift subsurface. In this proposed scenario the released H_2_, together with the available CO_2_ may trigger a dormant methanogenic community, resulting in the autotrophic production of methane, and thereby providing the basis for secondary, heterotrophic metabolic activity [19]. In addition, investigations of microbial composition and activity in mofette waters and mineral spring waters and surface sediments from the Cheb Basin have highlighted the role of acidophilic and methanogenic microbial processes in response to elevate levels of CO_2_, as well as the importance of sulfur and iron oxidizing and CO_2_ fixating microbial communities in these microhabitats [11-13, 15, 29, 30]. While these explorations provided some insights into how the unique geochemical conditions may shape microbial life across the Eger subsurface, to this date only one study has attempted to directly access and characterize microbial populations in Eger Rift subsurface sediments and rock formations. As part of a 2016 drilling campaign a 108.5 m deep core was recovered from the Hartusov mofette field and subsequently geochemically and microbiology analyzed [20, 22]. Analysis of the core lithology revealed Cenozoic sediments in the form of grey to brown and sandy to peaty mudstones, with lignite layers and root structures up to a depth of 90 mbs. Below 90 mbs core sections were characterized by weathered schists [22]. The recovery of CO_2_ rich sediments and the identification of a low permeable CO_2_-saturated saline aquifer around a depth of 80 mbs [22] further emphasized the significant role of CO_2_ in this ecosystem.

Microbial investigations targeted a 30 m section around the detected CO_2_ rich aquifer and resulted in the detection of a low biomass community, characterized by water and soil bacteria, specifically of the class Gammaproteobacteria [20]. Only minor signatures of microorganisms usually observed in acidic, high CO_2_ environments were detected, including low levels of methanogenic archaea and potentially autotrophic Comamonadaceae. The authors link the abundance and distribution of microorganism in the Eger Rift subsurface to frequently changing groundwater levels, as pump test data from the borehole suggest the presence of major fluid ascending channels, which indicate reoccurring vertical groundwater movement. Even though this drilling endeavor offered a valuable, first look into the microbial distribution and composition of Eger Rift subsurface sediments, the microbiological analyses solely focused on the area around the CO_2_ rich aquifer, effectively only providing a limited glimpse of what microbial life in this subsurface ecosystem may look like. To further extend the current understanding of the terrestrial biosphere in the Eger Rift and specifically evaluate the role of geologically derived compounds or substrates contributing to the development of microbial populations, additional efforts targeting broader regions of the Eger subsurface are needed.

In an effort to close this knowledge gap and advance the explorations of microbial life in the Eger Rift subsurface and other terrestrial subsurface regions, we evaluated drill core samples from a recent drilling campaign which reached a depth of 238 m. The main objective of this campaign was not just to provide a deeper and more comprehensive description of the microbiological composition, but to specifically evaluate the diversity and distribution of Archaea, as especially methanogenic Euryarchaeota may have the metabolic capability to utilize seismically released H_2_ in the presence of CO_2_. In addition, we hypothesized the Eger Rift subsurface to harbor microbial signatures contributing to the oxidation of inorganic compounds, including sulfur oxidizing and CO_2_ fixation microorganisms, and an overall microbial community shaped by the unique geological settings and elevated CO_2_ fluxes, including acidophilic and potentially halophilic taxa. Based on previous observations we also expected both the geochemical and microbiological composition of the recovered sediment and rock samples to be affected by the vertical groundwater movement and the existence of the described CO_2_ rich aquifer as well as similar structures, which may exist in deeper, previously uncharacterized regions of the Eger subsurface.

Using qPCR, 16S rRNA sequencing and fluorescent cell microscopy we were able to reconstruct microbial abundance and composition patterns across 26 samples, covering depths between 17 m and 230 m. Analysis of cation and anion concentrations across the core helped us to identify areas of increased ion dissolution, potential groundwater movement and CO_2_ aggregation, while microbial explorations provided novel insights into a fascinating microbial community highlighted by an unexcepted archaeal diversity. Findings from this study provide additional insights into the microbial structures found across the Eger Rift subsurface and advance the overall understanding of natural high CO_2_ subsurface ecosystems. Together with the data from previous Eger Rift explorations, our observations also provide the foundation for future efforts studying the interactions between volcanism, tectonics and microbiological activity in terrestrial subsurface environments with the goal to elucidate the impact of geological. processes on the deep biosphere.

## Materials and Methods

### Site description

Drilling of the F3 borehole was conducted in August 2019 at the Hartoušov Mofette field in the Cheb Basin in the Western part of the Eger Rift (Figure 1). The Hartoušov Mofette field and Cheb Basin in this area has been extensively described in previous work [22, 31-33], and is known for its unique patterns of CO_2_ degassing, with most heavy degassing observed in the central and northern part with emission rates of up 43 kg m^−2^ d^−1^ [34, 35]. The exact drilling position was selected in proximity to those of the previous two exploratory drilling campaigns (F1 and F2). The F1 drilling was conducted in 2007 [21, 36] and reached a depth of approximately 28 m below ground into a CO_2_-saturated, confined aquifer. The F2 borehole [36] was drilled in 2016 down to a depth ∼108 m. This drilling for the first time was conducted with the aim to evaluate whether the increased fluid and substrate flow can accelerate microbial life in active fault zones and CO_2_ conduits [20, 22].

**Figure 1:**
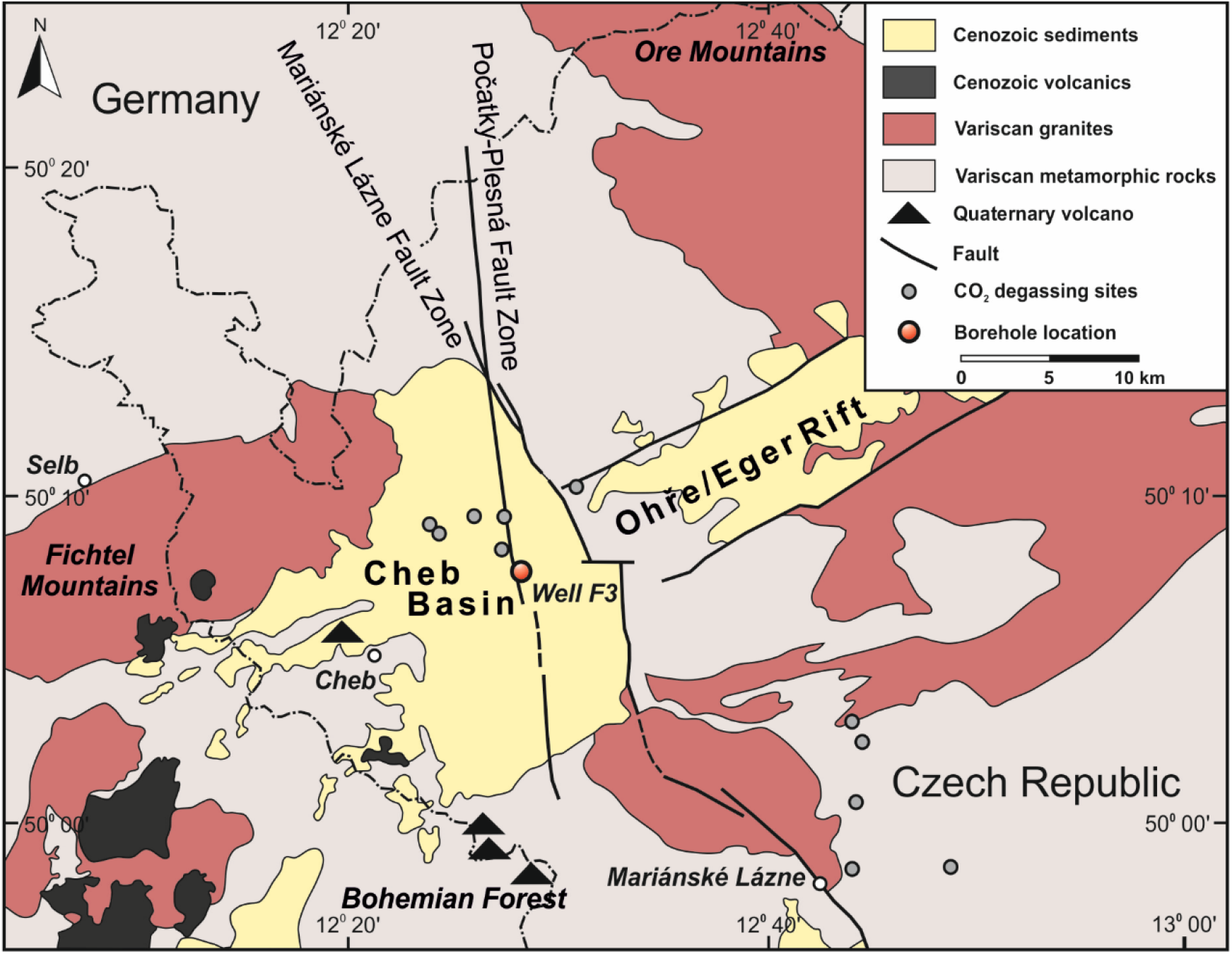
Geological map showing the location of the F3 borehole in the Cheb Basin in the western part of the Eger Rift (Czech Republic).

### Drilling and coring

Drilling of the 2019 drilling campaign started on August 9th 2019. First a bunker down to a depth 10 m was excavated and cemented. The upper 42m were then drilled using steel casing with an inner diameter of 85 mm (PQ). For this section core were retrieved in PVC liners of 1 m length and water was used as a drilling mud. Sample collection for geochemical and microbiological analysis started when a depth of 17 m was reached. From 40 m on drilling proceeded using HQ casing with an inner diameter for 63.5 mm (96 mm hole). Core sections were retrieved using 3 m aluminum liners. Drilling continued down to a depth of 238 m resulting in the recovery of 118 cores for total length of 163.15 m and a 82.3 % recovery rate. For the deeper drilling a mixture of bentonite and water was used for drill mud. To account for possible contamination of core material through drill mud, drilling was carried out under strict contamination control using fluorescent microbead tracer as reported previously [37, 38]. Tracer was added periodically to the drill mud and monitored to maintain a sufficiently high enough concentration. Drill mud samples were taken after core recovery, checked on site, and stored for later analysis (Table S1).

### On-site core subsampling and processing

After being brought up, core sections were immediately transferred to the mobile laboratory container of the GFZ Potsdam (BUGLAB) located in the vicinity of the drill site to allow core subsampling and sample processing under optimal conditions. Then 10 cm long whole round core sections were cut from the recovered cores. For downstream geochemical and culturing experiments core sections were then immediately placed in CO_2_ flushed gastight bags to preserve native conditions. Whole round core sections intended for microbiological analysis had the outer, drill mud contaminated rim removed on site (∽1cm), were placed in gastight bags and immediately frozen in liquid nitrogen. Geochemical and culturing samples were stored at 4°C throughout the remainder of the campaign and upon arrival at the GFZ German Research Center for Geosciences in Potsdam. Microbiological samples were stored in liquid nitrogen until arrival at GFZ Potsdam, where they were transferred to a -80°C freezer.

Remaining core sections were logged, cut into 1m pieces, labeled, stored in wooden core boxes and transported to the “Deutsche Bohrkernlager für kontinentale Forschungsbohrung” in Berlin Spandau for core description and long-term storage.

### Sample processing for genomic work

Frozen Core sections were slightly thawed and processed in a sterile clean bench (Thermo Scientific, Waltham, USA). To avoid collection of drill mud contaminated material core material was collected of the inner section of the whole round core. In addition, whole round core subsamples were taken for contamination control purposes. Up to 15 grams of the collected inner core sediment was stored in 50ml falcon tubes and frozen at -80°C until DNA extraction. As different DNA extraction approaches were employed up 1 gram core material was also stored in 2ml screw cap tubes at -80°C.

### Geochemical analysis

#### Ion leaching

The low pore water content of the collected core samples did not allow the collection of a sufficient amount of liquid. To assess the ionic composition samples underwent leaching (Blume et al., 2011). Briefly, 5 grams of dried sample were suspended in 25ml deionized water and agitated on a horizontal shaker for 90 minutes at 175 rpm. The samples were then centrifuged for 10 minutes at 175 rpm and the supernatant collected. The supernatant was filtered through a 0.45 μm filter and stored at 4°C. Duplicate samples from each core section were processed.

#### Ion Chromatography

Overall conductivity of the collected leachate was assessed using a TDS meter (WTW Multi 3420). Anion and cation composition of the leached pore water was measured using ion chromatography (IC) (Sykam Chromatography, Eresing, Germany). For anions the system consisted of a S5250 sample injector, a S150 IC-module, an anion separation column and a S3115 conductivity detector. The eluent was 6 mM Na_2_CO_3_ and 9 mM NaSCN. The eluent flow rate was set to 1 mL min^−1^ and the oven temperature was 50°C. Multi element standards from Sykam and Carl Roth were used. All samples were diluted 10X and measured in technical triplicates.

The cation chromatograph consisted of a S5300 sample injector, a ReproSil Cat separation column, and a S3115 conductivity detector. The flow rate was set 1.2 mL min ^−1^ eluent was 5mM H_2_SO_4_ and the oven was column oven was heated to 45°C. A 10-times diluted cation multi IC standard (Carl Roth) was used. Samples were measured undiluted as technical triplicates. IC chromatographs were analyzed using Chromstar7 software.

Differences in ionic composition across the sampled core were visualized by calculating a correlation matrix from z-score normalized ion concentrations and clustering on a PCA (principal component analysis) plot in Past4 [39].

#### Cell counts

For cell counts ∽3 grams of drill core sediment were collected in a 50ml Falcon tube and mixed with 2x volumes of 4% of filter-sterilized 2% (v/v) formaldehyde solution.

Cell extraction were then carried out based on work from Kallmeyer et al. 2008 [40] with some modification. For each fixed sample, 100-600 μL of homogenized slurry was mixed with 200 μL 2.5% (w/v) NaCl solution, 50 μL detergent mix (100 mM Na2EDTA•2H2O, 100 mM Na4P2O7•10H2O, 1% (v/v) Tween-80, autoclaved and filter-sterilized after cooling down) and 50 μL methanol. The mixture was vortexed for 30 min. 500 μL of filter-sterilized 50% (w/v) Nycodenz (Axis-Shield) solution was added from beneath the mixture through syringe needle (18-gauge). After centrifugation under 3000 ×g for 10 min, 800 μL of supernatant was collected to another tube. The remaining Nycodenz solution above the pellet was discarded, and 300 μL 2.5% (w/v) NaCl solution, 50 μL detergent mix as well as 50 μL methanol were added to the pellet. The mixture was sonicated in an ice-water bath for 4 × 10 s with 20 s between cycles at 25% power (BANDELIN SONOPULS, with ultrasonic probe MS73). Then a second time of Nycodenz density gradient centrifugation was performed on this μLtrasonicated mixture. The collected 800 μL supernatant was pooled with that from the first time and preserved for cell counting. To further dissolve the remaining clay-like particles in the supernatant, 1% (v/v) HF solution was added with a volume ratio of 1:1. The mixture was also undergone an ultrasonic bath for 15 min during the treatment of HF solution. The HF treated sample was diffused in 5 mL 0.5 M EDTA solution within 30 min to avoid excessive damage of HF on cells. The overall liquid was filtered through 0.2 um-pore-size filter membrane (GTBP, Millipore). 2 mL of detergent mix was also added before the filtration for a more even distribution of the sample on the surface of the filter membrane.

Cells were stained and mounted on slide with SYBR solution (300 μL VECTASHIELD mounting medium H-1000, 300 μL glycerol, 100 μL 1% (w/v) p-phenylenediamine, 10 μL commercial 10000* SYBR Green I stock, filled with H2O to 1 ml, filter-sterilized). Cells were counted under epi-fluorescence microscope (DM2000, Leica) with oil-immersion objective (HCX PL APO 100×/1.40, Leica) and light filter sets either Leica I3 or L5 [excitation filter of 470/40 nm and 480/40 nm (band-pass filter, center wavelength/bandwidth, same below) respectively, and suppression filter of 515 nm (long-pass filter, initiate wavelength) and 527/30 nm, respectively]. For each slide, at least 200 cells were counted unless 200 fields of view were screened.

### DNA extraction, clean up, and concentration

Due to the experiences of Liu et al. [20, 22] obtaining a sufficient amount of high-quality DNA from the recovered core sections was expected to be one of the major challenges of the study. To ensure optimal DNA recovery three different DNA extraction methods were employed.

First, DNA was extracted from up to 1 gram of sediment using a modified version of the Qiagen DNeasy PowerSoil kit (Qiagen, Venlo, Netherlands), as described previously [41, 42]. The major changes in comparison to the manufactures protocol were the addition of 10 mg/l lysozyme followed by an 1-hour incubation step at 50°C prior to bead beating. We also added three different types of zirconia beads instead of the kit supplied beads and used a 45 second bead beating step at 5 m/s instead of the protocol standard vertexing. The sample was extracted twice through bead beating to ensure maximum recovery (This means after centrifugation and removal of lysate post bead beating, another volume of extraction buffer was added and the bead beating, and centrifugation step was repeated). The remaining precipitation and DNA binding steps were carried out as described in the manufacturer’s protocol. Negative controls were included in every round of extraction.

DNA was also extracted using a modified Phenol Chloroform based method after Nercessian et al. [43]. Similar to the kit approach, 0.5 – 1.0 grams of sediment were treated with lysozyme and 800μl of an EDTA (0.5M) and phosphate (0.12M) based extraction buffer at 50°C for one hour followed by the addition of two sizes (0.1mm and 0.7mm) of zirconia beads and one size (3-4 mm) of glass beads. An equal volume of chloroform-isoamylalcohol and 10% SDS was added and the samples underwent bead beating for 45 seconds at 5 m/s. Samples were then centrifuged at 16,000 g for 10 minutes at 4°C. The lysate (supernatant) was collected and the bead beating step was repeated with another volume of extraction buffer. DNA was then precipitated by adding 0.5 volumes of isopropanol followed by incubation for one hour at room temperature. DNA was pelleted by centrifugation at 17000 g for one hour at 4°C. DNA pellets were washed with 70% ethanol and centrifuged at 17000 g for 10 minutes. DNA pellets were air dried and eluted in 100μl DNase free, PCR grade H_2_0. To achieve sufficient DNA extractions were performed in quadruplicates for each sample. Eluted DNA was then pooled and concentrated by precipitation with 100% Ethanol and 0.2M NaCl. Negative controls were included for each extraction run.

A third method used for DNA extraction in this study is based on a protocol by Rohland et al. [44]. First, 5.0 – 8.0 grams of core material were added to a 50ml falcon tube. Then 20ml of an EDTA based extraction Buffer were added (for 25ml: 22.5 ml 0.5M EDTA, 1.86 ml sterile PCR grade water, 12.5μl Tween 20, 10 625μl mg/ml Proteinase K). Samples were then incubated overnight (15-18 hours) at 37°C in a horizontally shaking incubator (80 rpm). Post incubation the supernatant was collected by centrifugation at 16,400 g for 2 minutes. Supernatant was then mixed with 2 volumes of binding buffer (5M guanidine hydrochloride, 40% (vol/vol) isopropanol and 0.05% (vol/vol) Tween 20). DNA was collected by running 5ml of the mixture over a Zymo silica spin column by centrifugation in a swing out rotor at 500 g for 5 minutes. The flow through was discarded and the step repeated until all of the mixture was run through the spin column. The spin column was transferred to a small 1.5ml low bind tube and DNA on the spin column was washed using an ethanol-based wash buffer (50% EtOH, 125 mM NaCl, 1mM EDTA). DNA was eluted in 80μl PCR grade H_2_0.

To limit bias through the applied DNA extraction protocol DNA from all three methods was pooled and cleaned using AMPure XP beads (Beckman Coulter, Pasadena, CA, USA). DNA concentrations were assessed using Qubit (Life Technologies, Carlsbad, CA, USA) and an Agilent tape station (Agilent, Santa Clara, CA, USA).

### Polymerase chain reaction (PCR) and Illumina sequencing

The V4 region of the 16S rRNA gene was amplified using universal 515F and 806R barcoded primers [45, 46] as described previously [20]. PCR reactions were run as quintuples for each sample for 30 cycles (30s at 95°C, 45s at 56°C, 60s at 72°C) to minimize the introduction of contamination. Extraction controls and template (PCR) controls were included in each PCR run using their own set of barcoded Primers. *E. coli* and microbial mock community DNA were included as positive controls. PCR products from the same samples were then pooled and cleaned using AMPure XP beads (Beckman Coulter, Pasadena, CA, USA). Concentration of PCR products was assessed using Qubit technology (Life Technologies, Carlsbad, CA, USA) and pooled in equimolar amounts. The pooled DNA library was concentrated with an Eppendorf Concentrator plus (Eppendorf AG, Hamburg, Germany) and sequenced on an Illumina Miseq Sequencer by a Eurofins Scientific SE.

### Bioinformatic and statistical analysis

Sequencing data was processed using the QIIME2 version 2019.10 [47]. Dual indexed sequencing reads were demultiplexed using CutAdapt [48]. DADA2 [49] was used to filter, denoise, and remove chimeras from the demultiplexed sequencing data. This included initial sequence truncations (250bp forward reads, 200 bp reverse reads). Quality-filtered paired end reads were then merged. All final sequences had a standardized read orientation and a minimum length of 200 bp. A sequence table was created resulting in Amplicon Sequence Variants (ASVs). The classify-sklearn [50] command was used to classify representative sequences identified through DADA2 using a pre-trained Naive Bayes classifier trained on Silva taxonomy database (v132) and assign taxonomic units (OTUs, 99%) [51, 52]. Singletons and OTUs assigned to chloroplasts and mitochondria from the OTU table. In addition, the resulting ASV table was manually screened and curated for contamination based on negative controls.

For illustration purposes taxonomic composition was summarized at the class and genus level. Class level taxonomy was visualized using the complex heatmap package in R [53]. The relative abundance of the most abundant 25 occurring ASVs was visualized on a bubbleplot using the phyloseq and ggplot2 packages in R [54-56].

The final curated ASV table was subsampled for a sequencing depth of 3,708 sequences (lowest sequence count) for alpha diversity analysis. Subsampling was conducted using the phyloseq package in R [54]. Microbial diversity was assessed by calculating observed OTUs, Shannon indices, and evenness indices using the estimate_richness command in phyloseq in R [54].

Inter sample diversity (beta-diversity) was determined by non-metric multidimensional scaling (NMDS) using Bray-Curtis dissimilarity distances in PAST4 [39]. The ASV table was Hellinger transformed for this analysis. Environmental parameters (here conductivity and ion concentrations) were z-score normalized fitted onto the ordination plot as vectors. Differences in microbial community structure among samples defined by environmental and physical parameters (depth, ionic strength) were assessed using analysis of similarity (ANOSIM) and permutational multivariate analysis of variance (PERMANOVA) calculations in Past4. Potential significant correlations between environmental parameters and biological data were examined and visualized using the cor function in R.

Co-occurrence network analysis was constructed using the Hsmic and Igraph packages in R. Correlations were determined by calculating Pearson correlation coefficients for all genera with a relative abundance of more than > 0.5% across all samples using the rcorr command in the package Hsmic [57]. Resulting correlation matrix was filtered for strength (R > 0.65) and significance (p < 0.05). Remaining correlations were turned into a network using the igraph package [58]. Modules based on the clustering were determined using the walktrap function and color coded using a custom RColorBrewer palette.

### Deposition of sequencing data

The Illumina MiSeq sequencing data of the generated 16S rRNA libraries was deposited at the European Nucleotide Archive (ENA). The dataset can be accessed at ENA or the National Center for Biotechnology Information (NCBI) under the accession number #PRJEB55581.

## Results

### Drilling and sampling

Drilling reached a final depth of 238m. Overall, 163.2 m (out of 198 m) of drill core were collected for a drill recovery rate of 82.3% and a total core loss of 17.7%. Core processing and sampling resulted in the collection of 55 samples, 28 for geochemical and 27 for biological analyses (Table S1). Two geochemical samples were compromised as the tubing broke during storage and transport. In addition, in two cases no microbiological samples could be taken, as the drill core material was crumbly and visibly contaminated by drill mud. Drill mud samples were collected periodically, and eight depths were selected as sequencing controls (Table S1). On-site observations suggested core sections below a depth of 50 meters to be frequently saturated with CO_2_, as the release of CO_2_ from was visible through bubbling and could be heard upon close examination.

### Core description

The core of well F3 can be divided into six lithostratigraphic units (Figure 2). The lowermost unit 1 from 239.5 to about 100.0 core meters consists of phyllitic mica schists and belongs to the Lower Paleozoic Saxothuringian basement of the Cheb Basin [59]. The light to medium grey phyllitic mica schists are predominantly fine-to medium-grained and show mostly moderately steeply dipping foliation in the form of schistosity and, in some cases, banding. They are mainly composed of quartz, feldspar and mica, mostly muscovite, and occasionally some chlorite. Pyrite occurs in crystals up to several mm in size, either distributed in mica schist or concentrated on planes of schistosity and mineralised joints. Bright bands and lenses, up to a few cm thick, consist of quartz and feldspar. The core interval at about 235 m contains a fracture plane with a dark-brown clay gouge and a surrounding fractured zone with numerous pyrite crystals several mm in size, indicative of a fault zone. In numerous core intervals, oblique or irregular pyrite-, quartz-or siderite-filled veins up to a few mm wide representing mineralized fractures cross the foliation and are predominantly siderite-filled in the upper core section of the unit. Particularly intense siderite mineralization in the form of both veins and irregular streaks and lenses is present in the core interval from ∽ 183.5 m to ∽ 152 m, with frequent occurrences of kaolinite. Irregularly distributed soft intervals are present in the phyllitic mica schists from ∽231 m depth upwards, increasing in frequency and length towards the top. The phyllitic mica schist is almost white and very soft in some intervals above ∽134 m, composed entirely of quartz, kaolinite and siderite, and showing poorly discernible foliation. At a depth of ∽ 113 m -∽ 113.5 m, irregular claystone clasts float in a matrix of kaolinite- and siderite-bearing phyllitic mica schists. Near the top of the unit, between ∽ 101 m and ∽ 100 m, three gypsum veins cross the mica schists. Minerals such as siderite, kaolinite and gypsum as well as a partial softening indicate that the phyllitic mica schist was heavily altered in some intervals. The kaolinite content, which tends to increase upwards, suggests that near-surface weathering processes in the Mesozoic and early Cenozoic [60] were most likely responsible for the kaolinitization. Siderite could have formed either from the influence of an overlying freshwater marsh during the Early Miocene [61] or from reduced conditions and increased pCO_2_ caused by CO_2_-rich deep crustal fluids. In any case, the occurrence of discrete core intervals with enhanced alteration indicates zones of increased permeability related to joints, schistosity, and faults.

**Figure 2:**
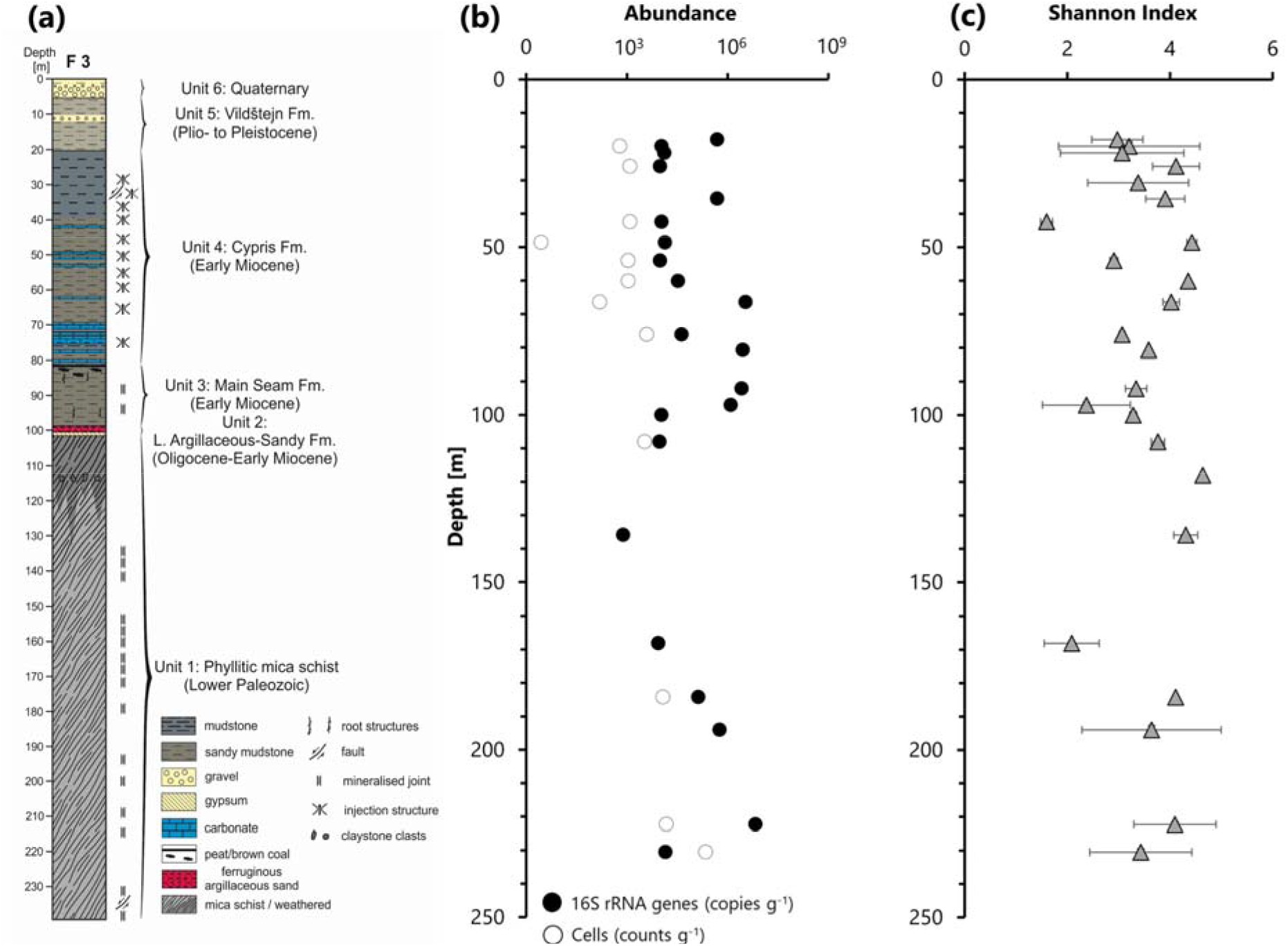
Stratigraphical and lithological description (a) of recovered Eger drill core, microbial abundance (b) measurements as determined by qPCR (16S rRNA gene copies) and fluorescent microscopy (cell counts), and diversity measurements depicted as Shannon Index (c).

The overlying unit 2 may only be present in the well from ∽100.0 m -∽99.65 m, but a core loss of almost 1.3 m does not allow for a definitive conclusion as to its thickness. It is a reddish colored, matrix-rich, clayey-silty sandstone that probably belongs to the Lower Argillaceous-Sandy Fm. [61]. The mineral content of quartz, kaolinite, siderite, hematite and some muscovite most probably reflect intense chemical weathering during deposition.

Unit 3 from ∽98.36 m to ∽80.50 m consists primarily of massive to weakly stratified gray to brown and frequently mottled sandy to peaty mudstone, which most likely represents the Early Miocene Main Seam Fm. [61]. The mineralogy consists of kaolinite, siderite, quartz and anatase as well as occasional greigite and crandallite. Several oblique to subvertical gypsum veins, a few mm thick, cross the sediments. Root structures occur primarily in the lower and upper intervals of the unit and together with fragments and layers of lignite in the uppermost interval indicate formation in a swamp environment.

Unit 4, ranging from approximately 80.5 to 20 m, consists predominantly of gray to green mudstones, often laminated. Primarily in the lower interval of the unit, sandy, partly peaty mudstones with plant remains are common, as well as carbonate layers, mainly dolomites, but also marl- and limestones. Ostracod shells are often concentrated on bedding planes. The mudstones are mainly composed of clay minerals such as muscovite/illite, kaolinite, smectite and mixed-layer minerals, as well as quartz, potassium feldspar, pyrite, greigite, zeolite, gypsum and analcime. Deformation structures are common, mainly sedimentary dikes, often surrounded by alteration zones. The laminated mudstones represent suspension deposits in a relatively deep lake, the lamination may reflect seasonal changes in the lake, while coarser sediments and detrital carbonates are likely turbidite deposits. The deformation structures seem to be mostly injection structures that could be the result of the ascent of CO_2_-rich deep crustal fluids. The unit most likely represents the Early Miocene Cypris Fm.

Unit 5, occurring between ∽20 m and ∽5.35 m, consists mainly of green to gray, moderately stratified to massive sandy clay. Gravel beds occur in the central section of the unit. This unit most probably represents lacustrine and alluvial sediments of the Pliocene to Pleistocene Vildštejn Formation [61].

The uppermost unit 6 from ∽5.35 m to the surface is mainly made up of moderately to poorly sorted clayey sand and gravel, which form fining-upward cycles of a few dm thickness. These are very probably Quaternary channel and floodplain deposits of the nearby Plesná River.

### Geochemical composition

The overall conductivity of the leaching products ranged between 98 μS cm^−1^ and 1726 μS cm^−1^ (Figure 3). Concentrations were relatively low across the first 70 m and then peaked in sediments from a depth around 75 m. Sediments from the intermediate section of the core (100 m – 200 m) were characterized by a lower ionic content, while below a depth of 200 m conductivity measurements increased again. To investigate the ionic distribution more closely, ion chromatography measurements for anions and cations were carried out.

**Figure 3:**
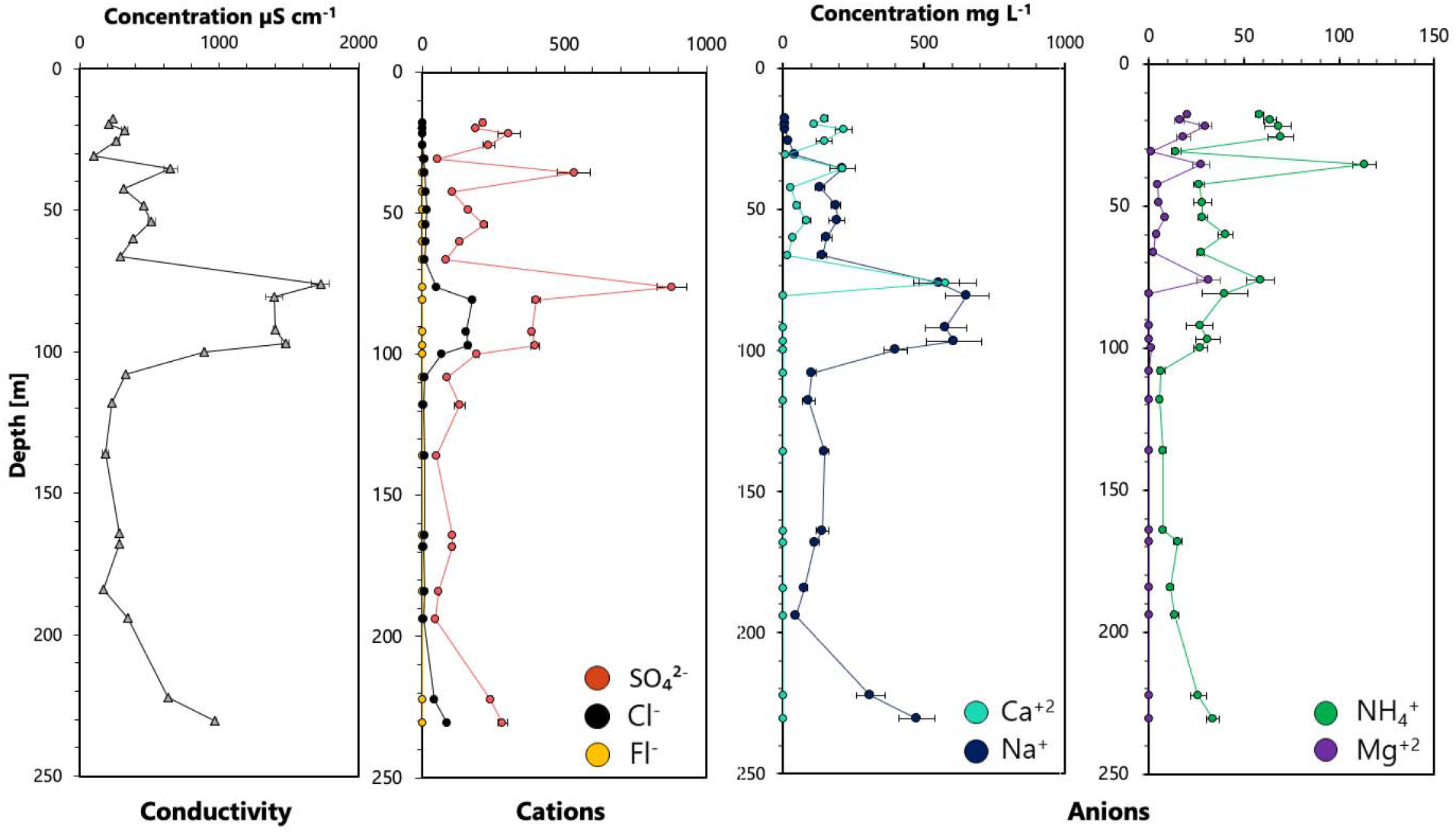
Geochemical composition, including overall conductivity, cation and anion concentrations across the recovered drill core profile.

Sulfate (up to 878.4 mg L^−1^) and Sodium (up to 653.6 mg L^−1^) were the most abundant ion species (Figure 3, Table S2). Levels were highest in sediments from depths between 80 and 100 m and increased again in rock samples from lower sections of the core. Chloride (up to 159.8 mg L^−1^) and Ammonium (up to 113.3 mg L^−1^) concentrations were lower overall, but followed the same pattern (Figure 3, Table S2). Calcium and Magnesium were only detected in sediments from the upper 76 m. Concentrations were elevated in the uppermost sediments (215.6 mg L^−1^ Calcium, 29.6 mg L^−1^ Magnesium) and then peaked at 76 m (576.0 mg L^−1^ Calcium, 31.2 mg L^−1^ Magnesium). Fluoride was barely detectable and was highest in sediments from around 30 m (2.3 mg L^−1^)

Principal component analysis (PCA) suggested ion composition to be associated with formation and thus depths, as especially samples from the main seam formation and deep phyllitc mica schist were found to cluster together (Figure S2).

### Microbial abundance and diversity

To assess the overall microbial abundance, distribution and diversity in Eger Rift sediments we employed cell counts via fluorescent microscopy, qPCR and 16S rRNA sequencing. Despite the difficult sample material, it was possible to recover DNA and prepare sequencing libraries from 24 different drill core sediment samples covering depths between 17 and 230 m (Table S3).

DNA extractions and 16S sequencing were performed in duplicate or triplicates for the majority of the samples (Table S3). Sequencing generated between 3,708 and 133,896 sequences (post trimming and contamination removal) per sample, with an average sequencing depth of 43,603 sequences (Table S3).

Quantitative PCR (qPCR) measurements confirmed the low biomass environment as measured values were just above the detection limit for most samples, ranging between 10^2^ and 10^6^ 16S rRNA gene copies per gram sediment extracted (Table S3, Figure 2). Highest values were detected between 60 and 100 m and in the deeper sediments recovered from a depth below 200 meters.

Cell counts using fluorescent microscopy were used to obtain an additional assessment of the overall biomass in the recovered drill core sediments but proved to be challenging. Cells could be counted in 13 samples and counts ranged between 10^1^ and 10^5^ cells g^−1^ (Table S3, Figure 2). Cell counts were highest in the deepest sediment below a depth of 185 meters.

Overall microbial diversity was assessed by determining Shannon alpha diversity indices as well as assessing the number of observed ASVs per 3,708 sequences. Highest diversities were observed around 50 m (414 ASVs, Shannon 4.3) and 120 m (576 ASVs, Shannon 4.6), while the least number of ASVs and the lowest Shannon index were detected at 42 m and 168 m (Figure 2). However, no significant trends or differences across samples could be identified.

### Inter sample diversity

Non-metric multidimensional scaling was conducted to evaluate potential differences in microbial community structure across the recovered drill core samples (Figure 4) and to compare recovered microbial communities to those identified in control drill mud samples (Figure S2). Drill mud samples were found to cluster separate from drill core samples, with only DM1 located in the vicinity of analyzed sediments (Figure S2).

**Figure 4:**
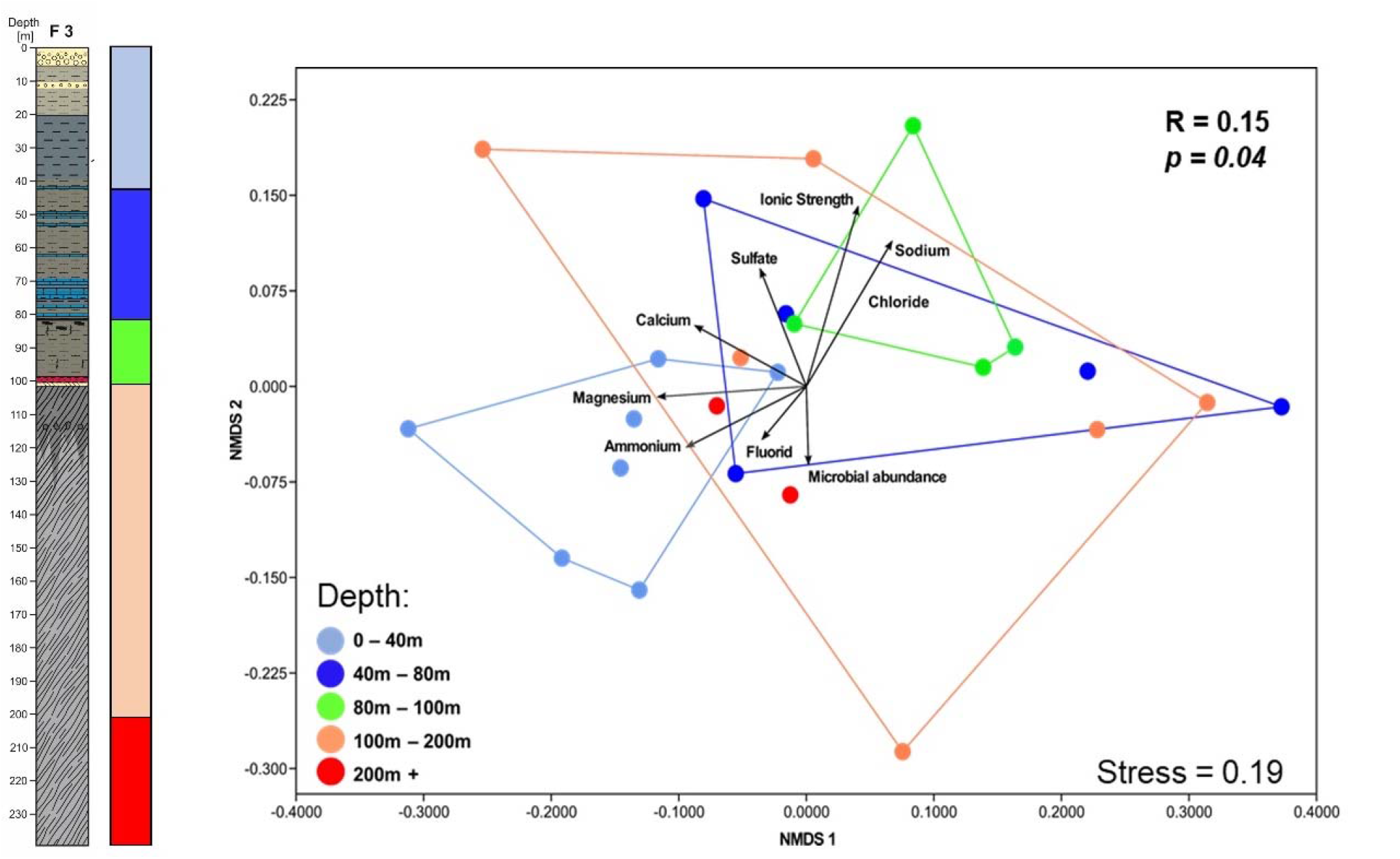
Non-metric dimensional scaling (NMDS) plot displaying microbial diversity across the recovered core samples. Samples are color coded by depth and have environmental parameters fitted as vectors.

Clustering by depth was observed for samples below 45 m and samples between 50 and 100 m, suggesting a shift in community across these regions. Microbial communities in deeper sediments were found to vary in diversity, as no distinct clusters could be identified. Similarly, samples associated with higher ion concentrations (ionic strength > 1000 mg L^−1^) were found to cluster close to each other, while the greatest diversity was observed among samples characterized by lower ion concentrations (Figure S3). Differences in community compositions in between samples grouped by depth or ionic concentration were further assessed using ANOSM (analysis of similarity) and PERMANOVA (permutational multivariate analysis of variance) calculations. These procedures further supported the graphical consideration, as differences between communities in shallow sediments (up to 50 m) and intermediate sediments (50 – 100 m) were found to be significant (ANOSIM, *p* = 0.02) or close to significant (PERMANOVA, *p* = 0.06). Similarly, grouping dependent on ion concentration could also be supported by this approach. Samples with high ionic content (>1000 ppm) were found to be microbiologically different from samples with very low (>200 ppm) and low (200 – 500 ppm) using both ANOSIM (*p* = 0.03 and *p* = 0.00) and PERMANOVA (both *p* = 0.05).

### Correlation and similarity analysis

To connect ecological, physical, and geochemical measurements a correlation analysis was conducted. Using spearman rank coefficients, it was possible to identify a strong, significant inverse correlation between sediment depth and the concentration of ammonium, calcium and magnesium ions (*p* > 0.05) (Figure S4). Weak, albeit insignificant, positive correlations between depth and the Shannon Index (*p* = 0.26) as well as the overall microbial abundance (based on 16S qPCR data) and the total ionic strength (R = 0.41, *p* = 0.12) could also be detected (Figure 5, Figure S4). Finally, we were able to detect a strong, significant correlation between the number of counted cells and depths (R = 0.73, *p* < 0.05), however as cells could only be counted for about half of the samples, we believe this finding should be construed with caution.

**Figure 5:**
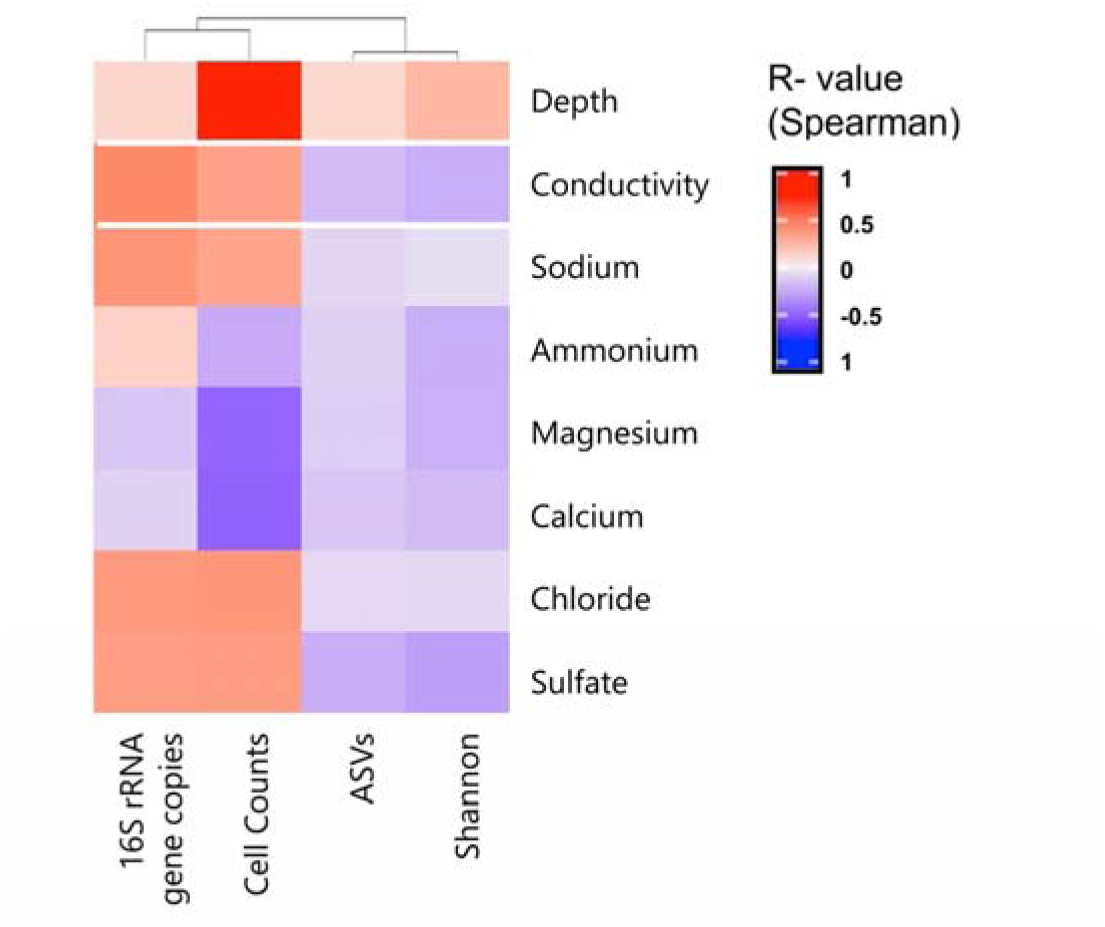
Correlation of microbiological parameters and environmental and geochemical parameters based on Spearman correlation coefficients.

### Microbial community structure

Evaluation of the microbial diversity and microbial composition across 230 m of drill core revealed a Bacteria dominated community enriched in soil and water associated microbes. Overall bacterial abundance varied between 100% and 90.6%, as Archaea were most abundantly detected in samples from depths between 30 and 42 m, (3.3 – 8.3%), 65 m (9.4%), 108 m (2.9%), and in deeper sediments between 185 and 230 m (1.0% -6.9%).

Generally, three different microbial community composition patterns could be observed based on the available 16S rRNA sequencing data. Down to a depth of 42 m drill core sediments were dominated by Alpha-und Gammaproteobacteria (Figures 6, S5), as especially the genera *Phyllobacterium* (up to 42.1 %), *Sphingomonas* (up to 43.9%), and *Pseudomonas* (up to 41.2%) were frequently detected. Phylogenetic classification of the most abundant *Sphingomonas* ASV suggested a close relationship to *Sphingomonas echinoides* strain 53234 and an environmental *Sphingomonas* clone previously identified in hydrocarbon contaminated soil (Figure 5, Figure S6). *Phyllobacterium* sequences were most closely associated with *Phyllobacterium bourgognense* and *Phyllobacterium zundukense*. The detected Pseudomonas ASVs related to *Pseudomonas peli* and were also highly similar to uncultured Pseudomonas sp. discovered in subsurface water [62]. Besides the occurrence of these abundant taxa, sediments above 42 m were also characterized by the frequent detection of the genera *Alshewenella* (up to 11.4%), *Caldiscericum* (up to 3.3%) and Chloroflexi of the taxon Dehalococcoida (up to 6.5%). Sediments recovered from the shallowest collected sample (17m) had a unique microbial signature as members of the Comamonadacaeae, particularly *Acidovorax* and (12.2%) and *Rhodoferax* (8.4%) were enriched.

**Figure 6:**
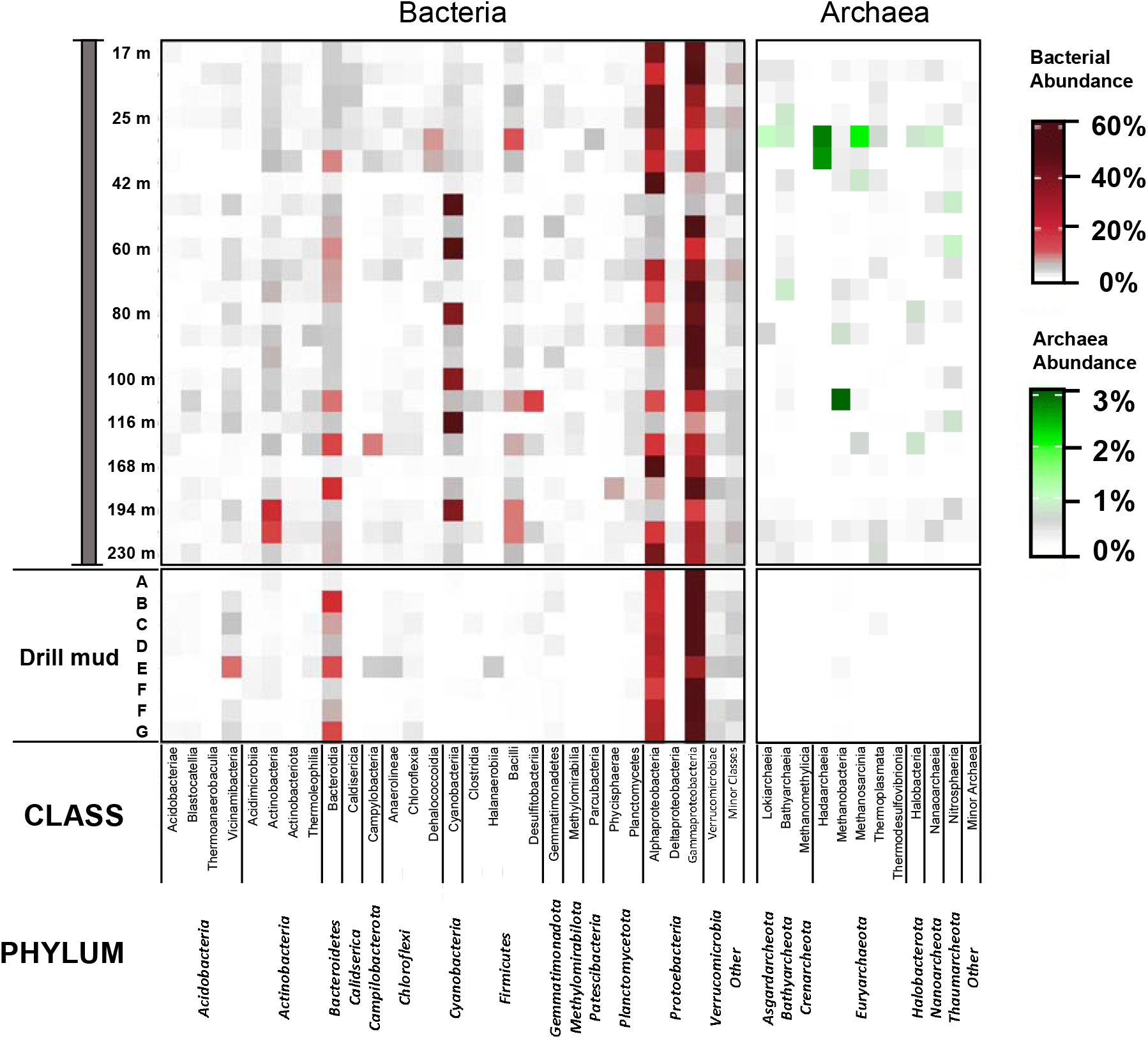
Class level microbial composition for Bacteria (red) and Archaea [63] across the recovered drill core samples and collected drill mud samples.

Below 42 mbs drill core microbial communities were more variable, as Alphaproteobacteria were less abundant and Gammaproteobacteria, especially *Pseudomonas* and *Alshewanella* abundance increased. Particularly notable was the frequent detection of several ASVs closely associated with freshwater Cyanobacteria species, especially *Cyanobium, Synechococcus*, and *Snowella*. However, high numbers of Cyanobacteria affiliated sequences were not detected continuously, but only at certain intervals (Figure 6, Figure S5, Figure S6). Cyanobacteria abundances were highest in samples from a depth of 48 m (69.4%), 60 m (53.3%), and 116 m (66.9%) (Figure 5, Figure S5, Figure S6). Between 108 m and 136 m Proteobacteria abundances were generally lower, as instead sediments were more enriched in sulfur cycle associated taxa such as *Sulfurimonas* (6.2% at 136 m) and the strict anaerobic, acidophilic *Desulfosporosinus* (15.1% at 108 m) as well as the chemolithoautrophic and halophilic taxon *Thiohalophilus* (4.5% at 135 m). Also notable was the discovery of *Roseisolibacter* (albeit at low abundances of up to 3.9% at 54 m and 87 m).

Deep sediments (150 m and below) were characterized by lower *Pseudomonas* counts, instead sequences affiliated with Firmicutes (*Bacillus*, 195 – 220 m) and Actinobacteria (*Streptomyces* 195 – 230 m) were detected more frequently. Another Cyanobacteria enrichment was identified at a depth of 195 m, as *Cyanobium* (31.2%) and Microcystis (4.1) signatures were enriched (Figure 6, Figure S5).

One specific focus of this study was the evaluation of the archaeal fraction, as the occurrence of CO_2_ and H_2_ utilizing, methanogenic Archaea is highly relevant to this type of degassing environment and may provide support for the microbial utilization of tectonically released substrates, specifically H_2_.

Analysis of the archaeal fraction revealed a large diversity, as 173 different archaeal ASVs could be detected. Our data indicates the presence of Euryarchaeota, Crenarchaeota and Nanoarchaeota throughout the drill core (Figure 7). Archaeal signatures were strongest in sediments from a depth around 30 to 40 m, as Hadarchaeia (up to 2.8%), Bathyarcheia (up to 4.5%), and *Methanosarcina* (up to 2.8%) were especially abundant. *Methanobacteria* sequences could be identified in most sediments, but were especially enriched at a depth of 108 mbs (2.7%). Thermophilic Thermoplasmata were also present in the majority of the evaluated core samples, and were most abundant at depth of 35 m (0.6%) and in the deepest sediments around 220 – 230 m (0.4 – 1.3%). Similar to the Thermoplasmata, Lokiarchaeia and Nanoarchaeia were detected at its highest frequencies (up to 0.9%) at these depths.

**Figure 7:**
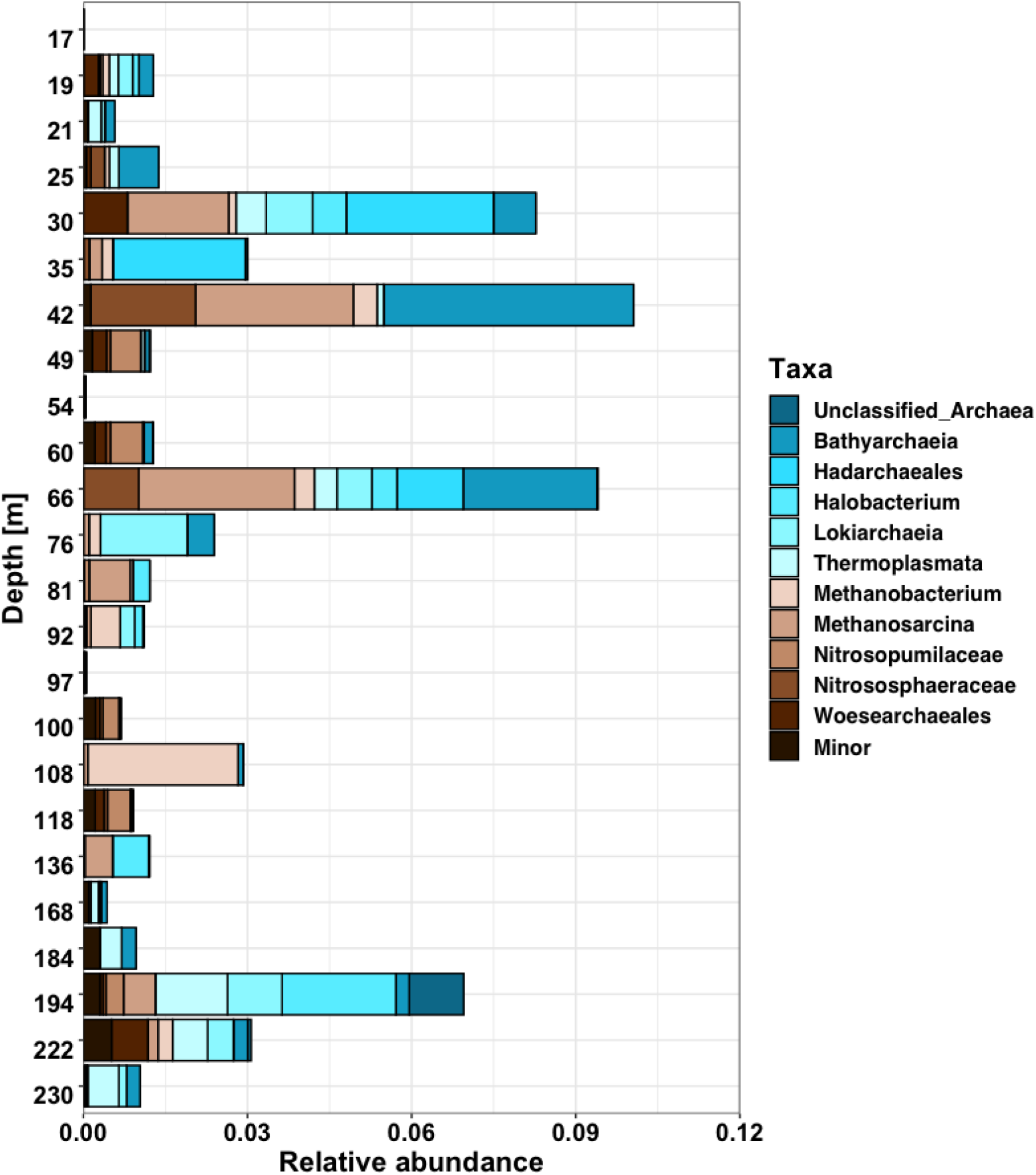
Distribution and relative abundance of Archaea across the recovered drill core sediments.

Potentially methanogenic taxa, especially *Methanosarcina, Methanobacteria* were detected frequently and found to be one of the most abundant groups, especially in sediments from an intermediate and shallow depths. Closer examination of the abundant *Methanosarcina* ASVs suggested the observed communities to be closely related to *Methanosarcina spelaei* strain MC-15, while detected *Methanobacteria* ASVs were closely affiliated with *Methanobactrium oryzae, Methanobacterium lacus* and unclassified *Methanobacterium* environmental sequences from lake sediments (Tree Figure S7).

Finally, several ASVs belonging to the ammonia oxidizing and often CO_2_ fixing Nitrososphaerota could be detected across the analyzed community (Figure 7). <u>Nitrosopumilaceae</u> (up to 0.6% at 60m) and Nitrososphaeraceae (up to 1.9% at 42m) were the most abundant and frequently occurring groups.

### Co-occurrence patterns

We used an advanced correlation analysis to identify microbial taxa co-occurring across the recovered drill core samples. The resulting network highlighted 17 different clusters, each representing one module (Figure 8, Table S4). Each vertex represents a genus., the most abundant and relevant to this ecosystem are listed in the adjacent table. The full list of taxa associated with each module is listed in table S4. Notable observations were the clustering of several Cyanobacterial taxa in Module 2, which also include the very small and often extreme environment associated Woesearchaeales and the subsurface associated Gaiellales.

**Figure 8:**
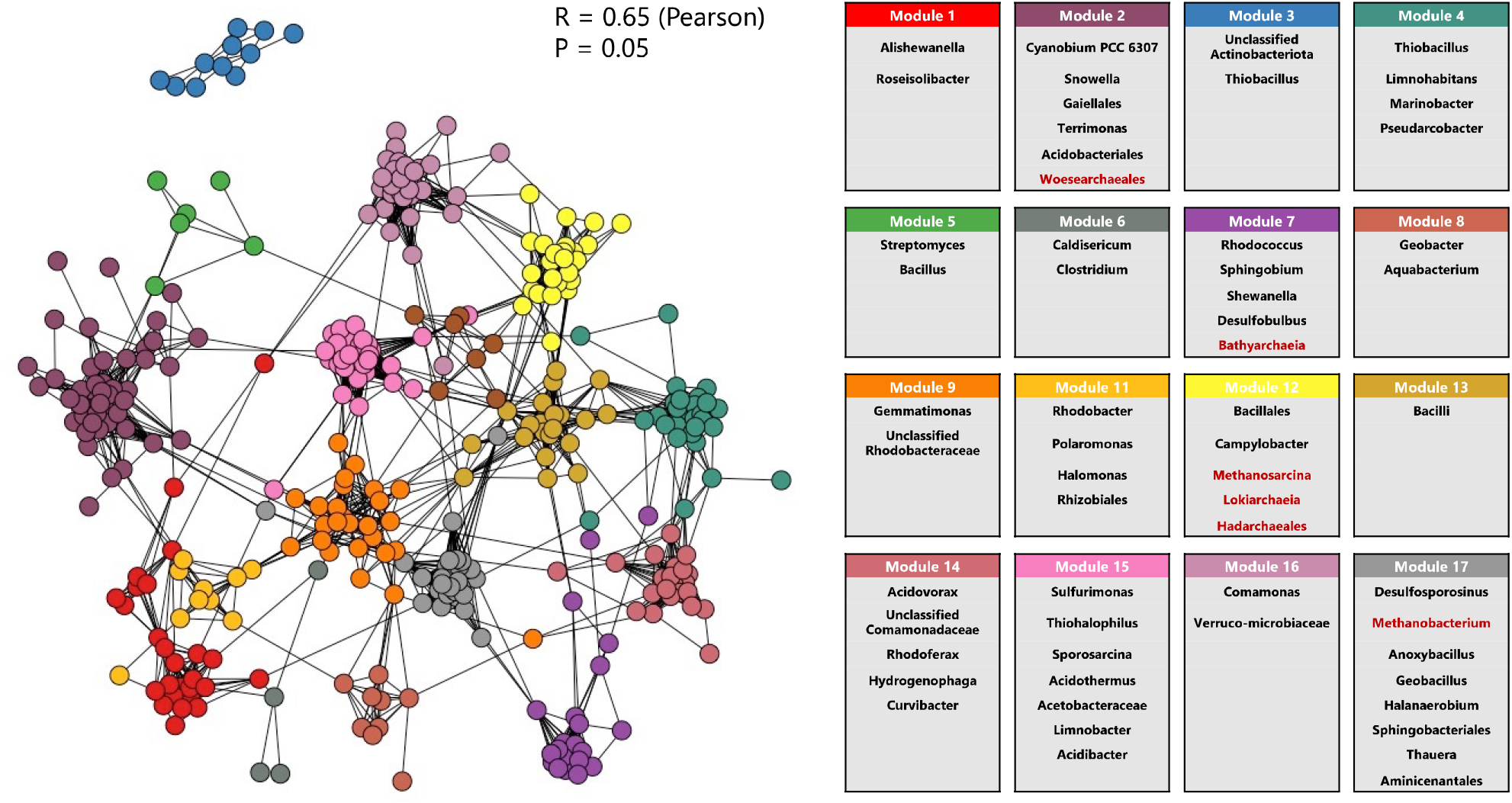
Co-occurrence network based on Pearson correlation (R > 0.65, p < 0.05) showing 17 different modules. Most abundant, identified bacterial and archaeal (red) genera associated with each module are listed in adjacent table.

In addition, the methanogenic taxa *Methanosarcina* (Module 12) and *Methanobacterium* (Module 17) were both found to co-occur with sulfate and iron reducers (*Desulfosporosinus* and *Geobacter*). Module 15 also included several acidophilic and sulfate utilizing taxa (*Sulfurimonas, Thiohalophilus, Acidothermus*), while Module 14 was especially characterized by the co-occurrence of taxa often in CO_2_ environment detected members of the Comamonadales (*Acidovorax, Rhodoferax, Curivbacter*).

## Discussion

### Unique ecosystem with abundant CO_2_

Explorations of terrestrial subsurface environments from a geochemical and especially biological standpoint offer the opportunity to study microbial life under unusual conditions, providing new insights into microbial distribution, diversity and metabolic capabilities, and can aid in the detection and description of bio-geo interactions.

As part of this study, we had the unique opportunity to investigate the geochemical and microbial composition of subsurface sediments from the Cheb Basin in the volcanically active Eger Rift region, obtained through the 238 m deep drilling into an active Mofette at the Hartoušov mofette field. Sample recovery and characterization supported the unique nature of the recovered core material, as sedimentology in the first 100 m changed frequently and allowed the identification of three distinct formations, including the Quaternary Vildstejn Formation (gypsum and sandy mudstone), the Cypris Formation (mudstone with frequent carbonate layers), and Main Seam Foundation (Sandy mudstone). Deeper drill core samples varied between slightly and heavily weathered schist, characteristic of recurrent groundwater movement and highlighted by the frequently present CO_2_ (CO_2_ bubbling could be observed when examining cores). This heterogenous section can therefore be considered its own system, with elevated transport through CO_2_ upwelling and groundwater fluctuations.

Despite difficulties arising from the various types of recovered core material and the overall low biomass in these samples, we were able to reconstruct and describe this unique microbial ecosystem through 16S rRNA amplicon sequencing. While superficially our data appears to emphasize the presence of a common soil and root microbial community, deeper investigation supports the existence of distinct microhabitats, shaped by the unique environmental, geological and geochemical conditions, and inhabited by distinct groups of chemoautotrophic, acidophilic and methanogenic microorganisms.

### Variable bacteria community dominated by soil and water microorganisms

Evaluation of the microbial community structure across the 230 m deep sediment and weathered schist samples highlighted a low biomass community especially abundant in heterotrophic soil and surface water bacteria. In agreement with the described sedimentology and core profile, communities seemed to vary within depth and sediment type across the upper 100 m, while deeper weathered schist populations were a lot more heterogeneous. These observations support earlier work by Liu et el. [20] from the same location, in which differences in community composition across formations at depths between 70 and 95 mbs were reported.

We found this section of the subsurface to be particularly enriched in microaerophilic or facultative anaerobic heterotrophs of the genera *Pseudomonas* and *Alshewanella*. While both these taxa have been identified in various subsurface settings, including hydrocarbon environments, drill cores and deep groundwater [41, 64-66], closer phylogenetic evaluation suggested these taxa to be closely related to strains associated with surface water bodies. Elevated concentration of anion (sulfate) and cations (sodium and calcium) in these sediment layers further emphasize this area to diverge from the surrounding subsurface. One potential explanation could be accumulation of CO_2_ in a subsurface aquifer, as suggested by Liu et al. [20] and Bussert et al. [22] or general groundwater movement in this area, which may also represent a link to surface water bodies, such as the river Pleśna, which runs in close proximity to the Hartousov mofette field.

Microbial communities in sediments from below 100 mbs, could not be distinguished by any geochemical or geological parameters, supporting the heterogeneous nature of the weathered schist, and the irregular/inconstant conditions predominating in this environment. We did observe an increase in cations in the deepest evaluated sediments, which correlated with a slightly higher microbial biomass, which could both be indicators for an additional CO_2_ rich aquifer at this depth. However, as only a small number of samples from these depths were available for this study, it is not possible to make any detailed statements on the microbial ecology in these regions.

### CO_2_ driven communities are not abundant but persistent

Our microbial ecology exploration did not reveal a rich microbial community of CO_2_ fixating microorganisms or even a strong abundance of acidophiles, as could have been expected based on the unique CO_2_ degassing conditions. Nevertheless, we identified taxa with these functional traits throughout the examined samples, suggesting this unique environmental feature to have somewhat of an impact on the ecology. Similar to previous studies from high CO_2_ environments [11, 67, 68], including the Eger subsurface [20, 32], we frequently identified taxa belonging to the Comamonadaceae family. While the Comamonadaceae contain a diverse group of water and soil bacteria, the here discovered genera *Acidovorax, Rhodoferax, Hydrogenphaga*, and *Curivbacter* have all been detected in high CO_2_ environments [12, 13, 69, 70], and have been suggested to fix or utilize CO_2_ or H_2_ in their likely mixotrophic lifestyle [11, 71]. Another finding supporting a somewhat CO_2_ driven bacterial community is the frequent detection of the taxa *Desulfosporosinus* and *Sulfurimonas*, both of which have been discovered in high CO_2_ and acidic environments [11, 13, 72-74]. Members of the taxon *Desulfosporosinus* are sulfate reducers, moderate acidophiles, and have the metabolic capability to fix CO_2_ into acetyl-CoA via the Wood-Ljungdahl pathway [75]. Similarly, *Sulfurimonas* are characterized by diverse energy metabolisms, which includes the potential to drive carbon fixation by rTCA cycling, via pathways of sulfur/hydrogen oxidation [76]. Additional, on-going work in our group also showed Eger sediment enrichments incubated under a H_2_/CO_2_ atmosphere to be highly abundant in *Desulfosporosinus* (Data not shown). Although we did not identify a correlation between the occurrence and abundance of this taxon with any of the geochemical parameters, specifically sulfate, these observations suggest this microbial group is able to thrive under high CO_2_ conditions and likely contributes to sulfur cycling in the Eger subsurface. Like *Sulfurimonas*, members of the single species genus *Thiohalophilus* are chemoautotrophic sulfur oxidizers. However, these organisms can assimilate CO_2_ via the Calvin–Benson–Bassham cycle [77] and usually occur in hypersaline lakes. While only detected occasionally in our samples, their appearance is also an indicator for a direct and indirect influence of CO_2_ in the Eger subsurface, as increased acidification can contribute to ion dissolution and may create areas of elevated salinity, suitable for these types of microbial populations.

Several other acidophilic taxa, including members of the Acidobacteriales and Acidothermales, could be detected throughout the Eger core samples, however only at limited abundances. These taxa were also previously detected in mofette and groundwater samples from the Cheb Basin [37,40]. Of interest was the co-occurrence of these taxa with the above-described acidophilic sulfur oxidizers *Sulfurimonas* and *Thiohalophilus*, as this further supports the concept of ecological niches, likely driven by CO_2_ and ascending saline groundwater and harboring acidophilic and halophilic microbial populations, existing throughout the Eger subsurface.

### The role of Cyanobacteria remains unclear

One of the most surprising findings of this study was the frequent detection of Cyanobacteria belonging to the phototrophic taxa *Cyanobium, Synechococcus, Microcystis*, as well as *Snowella* at very specific depth intervals. While the presence of Cyanobacteria in subsurface settings is not surprising in itself, as these types of microorganism have been detected in various types of subsurface environments at relatively high abundances previously [74, 78, 79], the fact that the here discovered taxa are usually associated with surface water bodies and an aerobic, phototrophic lifestyle [80] makes their discovery unexpected. Our initial thought was that these organisms may have been introduced through on-site or lab contamination. However, we included multiple drill mud and wet lab controls, which were also sequenced, and we could not find any of the abundant Cyanobacteria ASVs in these controls. Furthermore, the Cyanobacteria ASVs only occurred at very specific depth intervals, arguing against a general, external contamination source. Co-occurrence patterns also indicate these microbial populations to be native, as their presence was found to be statistically correlated to taxa typically found in the subsurface, including Gaiellales [81, 82], Woesearchaeota [83], and in soil and acidic environments, including *Terrimonas* and Acidobacteriales [29, 63, 84]. While at this point, we do not have a simple explanation for this phenomenon, the very distinct pockets of high cyanobacterial abundances offer a possible hypothesis. As discussed above, the Cheb Basin in the Eger region is home to numerous mineral water springs, which are connected to aquifers. Together with a subsurface environment that is characterized by frequent vertical groundwater movement, and the proximity of surface water bodies, like the adjacent river Pleśna, these attributes suggest the observed Cyanobacterial signatures may be washed seasonally or periodically into the here examined regions of the Eger subsurface. The fact that these organisms were not observed in samples from the 2016 drilling also supports their temporary presence. Going forward this hypothesis should be further investigated through periodical groundwater testing, or additional drilling efforts.

### Large diversity highlights importance of archaeal communities

One of the most important goals of this study, was to evaluate the presence and diversity of Archaea, as the Eger subsurface has been hypothesized to provide methanogenic substrates, H_2_ and CO_2_ through frequent swarm earthquake activity [18, 19, 22], with primary production through methanogenic archaea providing the basis for secondary heterotrophic lifestyle by bacteria. Previous studies have detected Archaea in Cheb Basin subsurface sediments [20] and mofette and spring waters from the region [32]. These efforts primarily detected Euryarchaea signatures, and especially in sediment samples archaea communities were low in abundance and diversity. Hence, it was hypothesized that seismic release of H_2_ triggers methanogenic activity, and that a small, dormant community of methanogenic archaea might become active following such an event. Our observations strongly support the importance of Archaea in the Eger subsurface and provide additional evidence that the terrestrial subsurface is home to a significant diversity of archaeal populations.

Similar to Liu et al. [20], we detected methanogenic Euryarcheota, as especially *Methanobacterium* and *Methanosarcina* signatures were identified across almost the entire length of the analyzed core. While relative abundances were highest in shallow and intermediate sediments down to 110 m, their presence in deeper samples suggest the potential for methanogenic activity to persist throughout. The genus *Methanosarcina* can produce methane using all three metabolic pathways for methanogenesis, and members of the *Methanobacterium* are specialized in the reduction of CO_2_ with H_2_ and are thus strict hydrogenotrophic methanogens [85-87]. Comparable to *Desulfosporosinus*, additional cultivation work in our group allowed us to grow *Methanobacterium* and *Methanosarcina* enrichments using CO_2_/H_2_ from various Eger sediment samples, even those where these organisms were present at minimal levels (Data not shown). The genus *Methanosarcina* is considered one of the most versatile microorganisms, as members belonging to this group can be acidophilic, thermophilic, or halophilic. Certain *Methanosarcina* species can also utilize acetate or methylated compounds to form CO_2_, but have also been suggested to produce acetate from carbon monoxide and thus provide the building blocks for secondary metabolisms, which may be essential in a subsurface environment otherwise scarce in organic substrates [8, 86-88]. Unfortunately, the here presented amplicon data does not allow us to specify which strain or species of *Methanobacterium* or *Methanosarcina* may be most abundant or specifically identify which type methanogenesis these organisms may be involved in. However, the frequent detection of these well-known methanogenic taxa provides additional evidence for the hypothesis of an ever-present Eger rift subsurface methanogen community. It is likely that once H_2_ become available as a substrate, methanogenic activity increases, producing biogenic methane signatures, which were observed by Brauer et al. [26, 27]. Both *Methanobacterium* and *Methanosarcina* ASVs were found to co-occur with sulfate and thiosulfate reducing and fermentative Proteobacteria and Firmicutes, such as *Desulfosporosinus, Halanaerobium* or *Bacillus* highlighting diverse microbial metabolisms. Generally, sulfate reduction is believed to restrict methanogenesis, questioning the active state of the here detected methanogens, however methanogenesis has been observed in zones dominated by sulfate reduction in various types of sediments [89, 90, 91]. In addition, methanogenesis and sulfate reduction can both occur when acetolactic, hydrogen producing, sulfate reducing bacteria and hydrogenotrophic methanogens coexist, which could be the case in the Eger subsurface.

While the detection of methanogenic archaea provided new insights on methanogenic processes in the Eger rift subsurface, our exploration also highlighted various other Euryarchaeota species. Of special interest were the frequent detection of acidophilic Themoplasmata and halophilic Halobacteria [87, 92], whose ubiquity can most likely be attributed to their surroundings and thus may be additional examples of microbial populations established and driven by the unique CO_2_ degassing conditions in the Eger Rift subsurface. Another striking finding was the high representation of Hadarchaeales sequences in several samples, as this relatively newly established group of *Candidatus* microorganisms is closely associated with both the marine and terrestrial subsurface settings and was first discovered in acidic hot springs [8, 93]. Phylogenetically, Hadarchaeales have been classified as Euryarchaeota, and are known for their versatile metabolism, which has been described to include the capability to oxidize hydrogen and one carbon compounds [76, 94]. Further evidence for a diverse and thriving archaeal community in the Eger subsurface were the detection of several Proteoarcheota taxa. Bathyarcheaota and Nitrosphaera signatures were detected throughout almost all samples and especially enriched in intermediate sediments, while Lokiarchaeia could be detected in shallow and deep samples. Bathyarchaeota are known to be a globally occurring, sedimentary archaeal phylum, which is found in various types of environments, including marine and terrestrial sediments, hot springs, and hydrothermal vents [30, 95-98]. While their direct role in methanogenesis is not clear, certain Bathyarchaeia strains have the metabolic capability to utilize CO_2_ through reductive acetogenesis via an archaeal variant of the Wood–Ljungdahl pathway [99] and interact with methanogenic communities [95], suggesting their frequent detection may be tied to the presence of CO_2_ and H_2_ in the Eger subsurface.

Similar to Bathyarchaeota, Nitrosphaera are known for their ubiquitous global distribution, occurring primarily in soils, but having been detected in various environments, including high salinities, low pH settings (down to 3.5) and hyperthermal environments [100-102]. Metabolically this group of microorganisms has the ability to oxidize ammonia aerobically or anaerobically and thus sustain autotrophic growth [103]. Interestingly, Nitrosphaera was found to correlate with overall microbial diversity moderately, but significantly, and several genera (Nitrosopumilaceae, *Nitrosarchaeum*) were found to co-occur with bacterial nitrite and ammonia oxidizers, as well Cyanobacteria, suggesting their distribution could also be influenced by the surrounding surface water and even agricultural run-off. Lokiarchaeia are another example of recently discovered Candidatus phyla and belong to the Asgard superphylum. Originally detected in mid ocean sediment samples, closely related genomes have since been detected in various terrestrial anaerobic and microaerophilic habitats, including hot springs, groundwater, and cave systems [104, 105]. Lokiarchaeia have been suggested to share a common ancestor with Eukaryotes, as shared protein signatures could be identified, however very little is to this point known about their metabolic capabilities and lifestyle.

Altogether our exploration into the archaeal subsurface communities highlight a diverse adapted community, shaped by the unique geological and environmental features. Our observations also provide additional evidence that methanogenic archaea are likely key contributors to microbial processes in the Eger rift subsurface and should have the ability to utilize the available resources, CO_2_ and when available H_2_. Nevertheless, it needs to be stated that the here presented results are based on relatively short 16S rRNA amplicon sequencing data, which only allow classification to a certain taxonomic level and do not provide any information on viability or metabolic capability. Additional analyses are needed to build on the presented data. On-going work in our group is currently further examining the metabolic role of archaea through enrichments and metagenomic explorations. Initial experiments have been successful, measuring methane production, and allowed us to reconstruct nearly complete draft genomes for *Methanobacterium* and *Methanosarcina*, which are currently being analyzed. Metagenomic data may also help to obtain additional insights on various unculturable archaeal groups, so we have intensified our efforts to obtain enough DNA from these difficult to work with, low biomass samples.

### Biosphere summary and study limitations

Evaluation of the microbial composition in Eger rift drill core samples highlighted the need to look beyond first impressions and displayed a not highly abundant, but distinct and consistent community driven by the unique geological and environmental features found across the evaluated core profile, such as increased pH, elevated salinity, CO_2_ degassing and frequent groundwater movement. While overall low detected biomass further negates the hypothesis of a “microbial hotspot”, our data also emphasizes the potential for a variety of microbial processes and heterotrophic, autotrophic and chemolithotrophic lifestyles. Besides the strong presence of soil and water associated Proteobacteria, such as *Pseudomonas*, which are certainly found in many similar environments and are known for their adaptability and metabolic flexibility, especially the high archaeal diversity stands out as scientifically compelling. While limited archaeal populations were detected in previous Eger microbiological studies and are certainly seen in other terrestrial subsurface environments, the here observed variety and even abundance in distinct samples emphasizes their perseverance and importance in extreme and especially CO_2_ rich settings and the deep biosphere overall.

Although only a limited number of geochemical measurements could be taken, our analysis showed that the distribution of ions is related to the microbial community, as elevated conductivity and increases in certain ionic species coincided with changes in microbial composition. Nonetheless, it is unclear whether both ion distribution and microbial community are driven by the environment, or if the one directly impacts the other. As dissolution of ions increases salinity and thus creates an environment more favorable for halophilic organisms, we expected to identify correlations between specific taxonomic groups and the abundance of ionic groups, none of which were observed. It also needs to be noted, that there certainly are additional environmental and geochemical parameters, which may impact the microbial community, which could not be included in this study.

## Conclusions

The terrestrial subsurface has become an area of significant scientific, industrial and societal importance, providing resources like groundwater, minerals, metals and hydrocarbons. Despite the growing relevance, subsurface microbial life and diversity, especially considering the various types of unique subsurface settings existing around the globe, remains limited. Additional investigations targeting both microbial communities and processes and their interactions with the surrounding geological and geochemical characteristics are needed to explore microbial behavior and address questions regarding metabolic potential and community adaption. Here, we had the unique opportunity to study microbial life in one of the world’s most distinct subsurface system, the area below the Hartoušov Mofette field in the Cheb Basin of the Eger Rift, an area abundant in sediments and known for frequent swarm earthquake activity and steady mantle derived CO_2_ degassing. Through our investigation, we were able to characterize the ionic composition, microbial abundance, and microbial community structure across a 238 m drill core. These analyses provide novel data on how high CO_2_ concentrations in the subsurface may impact geological structures and shape microbial behavior. Overall, our data suggests large parts of geochemical and microbial to be driven by groundwater movement, CO_2_ upwelling and the accumulation of CO_2_ in aquifers, as samples from previously not analyzed regions below 100 m were highly heterogeneous and characterized by varying types of microorganisms, with the capability to pursue aerobic, anaerobic, heterotrophic, autotrophic and chemolithotrophic lifestyles. A deeper look revealed a surprisingly diverse community of Archaea, providing strong support that both methanogenic as well as autotrophic and acidophilic archaeal communities reside in the Eger subsurface, likely taking advantage of substrates released through the volcanic properties of the region, such as H_2_, and the omnipresent CO_2_. The discovery of abundant Cyanobacteria signatures at distinct depths emphasizes the Eger Rift’s unique subsurface features and shows that additional and temporal explorations are going to be necessary to shed more light on this phenomenon. At this point, our findings suggest that certain subsurface areas represent frequently changing environments, impacted by varying groundwater levels, run-off from the surrounding surface and are in potentially regular exchange with surrounding surface water bodies. While the here introduced data provides novel insights into the geochemical and microbiological composition of the Eger Rift subsurface, especially for the deeper regions, we believe the next steps are to obtain greater taxonomic resolution, study microbial behavior, and evaluate microbial processes in more detail. These efforts are on their way as we are already in the process of employing metagenomics and growing enrichments to study the Eger Rift subsurface biosphere in more detail. Going forward, on-going and future investigations, together with data from this study, will provide additional, valuable insights on microbial life in one of the most unique subsurface environments on earth, and help scientists study bio-geo interactions in CO_2_ enriched deep biosphere settings.

## Funding and Acknowledgements

The ICDP-Eger Rift project was funded by the International Continental Drilling Program, the German Science Foundation (DFG) -project nos. 419880416, 419459207 and 419909358, the Czech Science Foundation, GFZ Potsdam and large project of Czech infrastructure CzechGeo (CZ.02.1.01/0.0/0.0/16_013/0001800). We also received co-funding from the Swedish National Research infrastructure for scientific drilling Riksriggen at Lund University, Sweden.

