## Supplemental Material for "Microbial diversity and community dynamics in an active, high CO_2_ subsurface rift ecosystem"

Supplemental Information:

| **Table S1:** Summary of collected Eger drill core samples | | | | | | |
| --- | --- | --- | --- | --- | --- | --- |
| **Sample**  **ID** | **Depth**  **[m]** | **GEOCHEM,**  **CO_2_ flushed,**  **4^o^C** | **MBIO,**  **Liquid N_2_ flash freeze** | **Drill mud sample Seq** | **Cont. Control** | **Comment** |
| 17 | 17.8 | X | X |  | X | In plastic liner |
| 19 | 19.8 | X | X |  | X | In plastic liner |
| 21 | 21.8 | X | X |  |  | In plastic liner |
| 25 | 25.8 | X | X |  | X | In plastic liner |
| 30 | 30.7 | X | X |  | X | In plastic liner |
| 35 | 35.5 | X | X |  | X | In plastic liner |
| 42 | 42.2 | X | X | X | X | In plastic liner |
| 44 | 48.6 | X | X |  |  |  |
| 46 | 54 | X | X |  | X |  |
| 48 | 60.1 | X | X |  |  |  |
| 50 | 66.4 | X | X | X | X |  |
| 52 | 71.1 |  |  |  |  | Fractures and loose material, no MBIO sample, GEOCHEM sample broke |
| 54 | 76.1 | X | X | X | X |  |
| 56 | 80.7 | X | X |  | X |  |
| 58 | 88.4 | X |  |  |  | Sample compromised |
| 59 | 92.1 | X | X |  | X |  |
| 61 | 97 | X | X |  |  |  |
| 62 | 100 | X | X | X | X |  |
| 65 | 118.9 | X | X |  | X |  |
| 68 | 135.9 | X | X |  |  |  |
| 74 | 136.1 | X | X |  |  |  |
| 80 | 154 | X | X | X |  | No PCR product |
| 86 | 168.1 | X | X | X | X |  |
| 93 | 184.2 | X | X | X |  |  |
| 98 | 194 | X | X |  | X |  |
| 99 | 195.2 | X | X |  | X | Contaminated |
| 112 | 222.3 | X | X | X | X |  |
| 116 | 230.5 | X | X |  | X |  |

| **Table S1:** Geochemical and features of the recovered Eger drill core sediment samples | | | | | | | | | | |
| --- | --- | --- | --- | --- | --- | --- | --- | --- | --- | --- |
| **Sample ID** | **Depth [m]** | **Formation** | **Ionic Strength*** | **Sodium** | **Ammonium** | **Magnesium** | **Calcium** | **Fluoride** | **Chloride** | **Sulfate** |
|  |  |  | **µS** | **mg/L** | | | | | | |
| 17 | 17.8 | Quaternary Vildsteijn Fm. | 241 | 7.72 | 57.86 | 19.96 | 147.99 | 0.00 | 0.71 | 212.34 |
| 19 | 19.8 | Quaternary Vildsteijn Fm. | 208 | 6.84 | 63.55 | 16.02 | 111.67 | 0.00 | 0.41 | 185.80 |
| 21 | 21.8 | Cypris Fm | 321 | 7.89 | 67.89 | 29.62 | 215.61 | 0.00 | 0.48 | 303.98 |
| 25 | 25.8 | Cypris Fm | 263 | 18.97 | 69.15 | 18.06 | 145.95 | 0.32 | 0.90 | 233.68 |
| 30 | 30.7 | Cypris Fm | 98 | 41.33 | 14.16 | 0.89 | 6.65 | 2.34 | 5.88 | 54.02 |
| 35 | 35.5 | Cypris Fm | 652 | 210.09 | 113.29 | 27.22 | 210.89 | 0.00 | 7.99 | 533.05 |
| 42 | 42.4 | Cypris Fm | 316 | 132.21 | 26.19 | 4.27 | 25.73 | 0.44 | 10.79 | 103.76 |
| 44 | 48.6 | Cypris Fm | 455 | 187.36 | 28.11 | 5.31 | 50.53 | 0.34 | 15.28 | 161.16 |
| 46 | 54 | Cypris Fm | 515 | 191.10 | 28.12 | 8.24 | 84.43 | 0.18 | 11.43 | 218.79 |
| 48 | 60.1 | Cypris Fm | 383 | 155.73 | 40.05 | 3.67 | 33.40 | 0.34 | 9.94 | 131.84 |
| 50 | 66.4 | Cypris Fm | 295 | 139.18 | 27.11 | 2.09 | 17.47 | 0.45 | 8.02 | 83.91 |
| 54 | 76.1 | Cypris Fm | 1726 | 555.22 | 58.70 | 31.19 | 574.99 | 0.00 | 48.22 | 878.36 |
| 56 | 80.7 | Main Seam Fm | 1394 | 653.62 | 39.79 | 0.00 | 0.00 | 0.95 | 176.42 | 400.41 |
| 59 | 92.1 | Main Seam Fm | 1399 | 577.28 | 26.62 | 0.00 | 0.00 | 0.72 | 153.52 | 385.13 |
| 61 | 97 | Main Seam Fm | 1475 | 606.99 | 30.98 | 0.00 | 0.00 | 0.73 | 159.75 | 397.83 |
| 62 | 100 | Phyllitic mica schist | 892 | 400.04 | 26.95 | 1.39 | 0.00 | 0.66 | 66.93 | 191.17 |
| 65 | 108 | Phyllitic mica schist | 329 | 102.88 | 6.32 | 0.00 | 0.00 | 0.20 | 7.13 | 85.96 |
| 68 | 118 | Phyllitic mica schist | 228 | 90.78 | 5.49 | 0.00 | 0.00 | 0.00 | 4.44 | 129.81 |
| 74 | 135.9 | Phyllitic mica schist | 184 | 149.08 | 7.38 | 0.00 | 0.00 | 0.22 | 8.59 | 48.01 |
| 80 | 164 | Phyllitic mica schist |  | 138.69 | 7.51 | 0.00 | 0.00 | 0.00 | 6.68 | 104.39 |
| 86 | 168.1 | Phyllitic mica schist | 282 | 114.56 | 15.27 | 0.00 | 0.00 | 0.17 | 5.45 | 105.37 |
| 93 | 184.2 | Phyllitic mica schist | 169 | 77.04 | 11.22 | 0.00 | 0.00 | 0.24 | 7.38 | 58.09 |
| 98 | 194 | Phyllitic mica schist | 346 | 46.98 | 13.66 | 0.00 | 0.00 | 0.15 | 4.71 | 45.04 |
| 112 | 222.3 | Phyllitic mica schist | 630 | 311.20 | 25.90 | 0.00 | 0.00 | 0.25 | 42.19 | 238.22 |
| 116 | 230.5 | Phyllitic mica schist | 970 | 473.46 | 33.45 | 0.00 | 0.00 | 0.36 | 84.43 | 282.02 |
| *As assessed with Probe | |  |  |  |  |  |  |  |  |  |

| **Table S2:** Microbiological features of recovered and analyzed Eger drill core sediments | | | | | | | |
| --- | --- | --- | --- | --- | --- | --- | --- |
| **Sample ID** | **Depth [m]** | **Generated 16S rRNA sequences*** | **qPCR** | **Cell counts** | **# ASVs*** | **Shannon*** | **Evenness*** |
|  |  |  | **16S rRNA gene copies per gram** | **Cells per gram** | **rarified to 3473 sequences** | | |
| 17 | 17.8 | 171008 | 4.89E+05 | N/A | 185 | 2.97 | 0.11 |
| 19 | 19.8 | 44430 | 1.07E+04 | 6.14E+02 | 220 | 3.20 | 0.21 |
| 21 | 21.8 | 23870 | 1.30E+04 | N/A | 135 | 3.07 | 0.20 |
| 25 | 25.8 | 45415 | 9.63E+03 | 1.23E+03 | 318 | 4.12 | 0.24 |
| 30 | 30.7 | 9815 | N/A | N/A | 109 | 3.38 | 0.34 |
| 35 | 35.5 | 10549 | 4.89E+05 | N/A | 101 | 3.91 | 0.50 |
| 42 | 42.4 | 33042 | 1.07E+04 | 1.23E+03 | 71 | 1.60 | 0.07 |
| 44 | 48.6 | 73047 | 1.35E+04 | 2.79E+00 | 415 | 4.43 | 0.20 |
| 46 | 54 | 89002 | 9.63E+03 | 1.08E+03 | 253 | 2.91 | 0.07 |
| 48 | 60.1 | 38256 | 3.31E+04 | 1.09E+03 | 406 | 4.35 | 0.19 |
| 50 | 66.4 | 26448 | 3.41E+06 | 1.54E+02 | 194 | 4.02 | 0.29 |
| 54 | 76.1 | 10692 | 4.22E+04 | 3.89E+03 | 100 | 3.07 | 0.24 |
| 56 | 80.7 | 3702 | 2.83E+06 | N/A | 140 | 3.58 | 0.26 |
| 59 | 92.1 | 664882 | 2.61E+06 | N/A | 174 | 3.34 | 0.17 |
| 61 | 97 | 79349 | 1.23E+06 | N/A | 182 | 2.37 | 0.07 |
| 62 | 100 | 52352 | 1.06E+04 | N/A | 276 | 3.28 | 0.12 |
| 65 | 108 | 5102 | 9.30E+03 | 3.38E+03 | 113 | 3.76 | 0.41 |
| 68 | 118 | 113156 | N/A | N/A | 576 | 4.64 | 0.18 |
| 74 | 135.9 | 42524 | 7.67E+02 | N/A | 342 | 4.31 | 0.22 |
| 86 | 168.1 | 38711 | 8.54E+03 | N/A | 144 | 2.09 | 0.06 |
| 93 | 184.2 | 14641 | 1.33E+05 | 1.18E+04 | 158 | 4.11 | 0.39 |
| 98 | 194 | 62632 | 5.72E+05 | N/A | 210 | 3.64 | 0.18 |
| 112 | 222.3 | 25142 | 6.75E+06 | 1.49E+04 | 217 | 4.09 | 0.38 |
| 116 | 230.5 | 49697 | 1.40E+04 | 2.21E+05 | 171 | 3.43 | 0.25 |
| * average from up to three replicates | | |  |  |  |  |  |

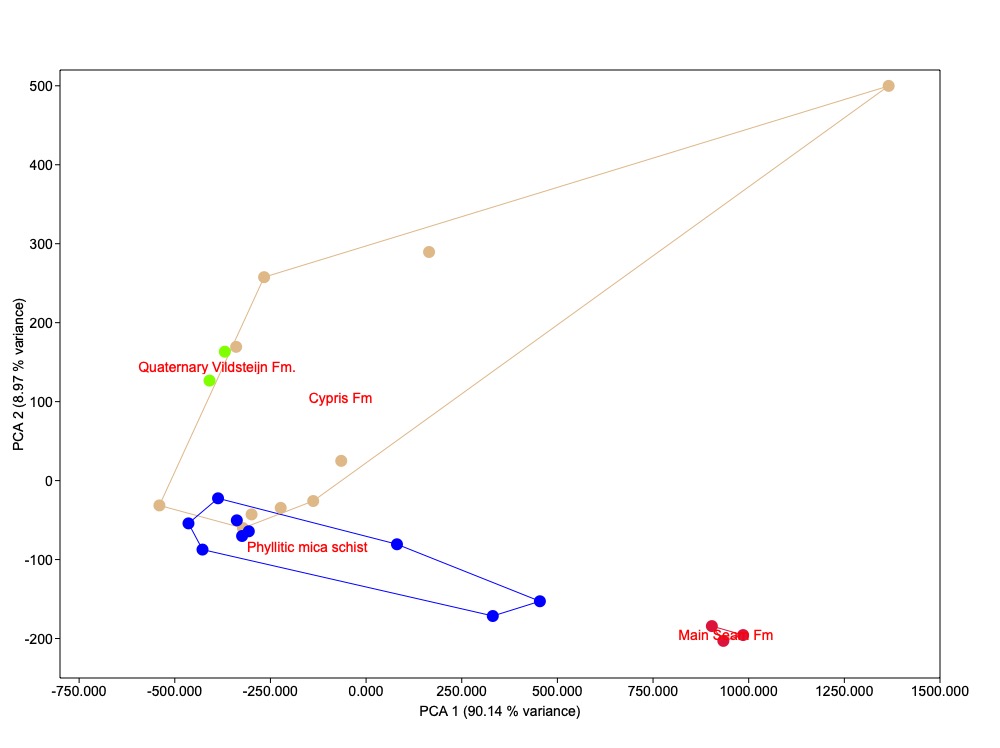

**Figure S1:** PCA plot based on ionic composition of recovered from Eger Rift sediments and color coded by formation.

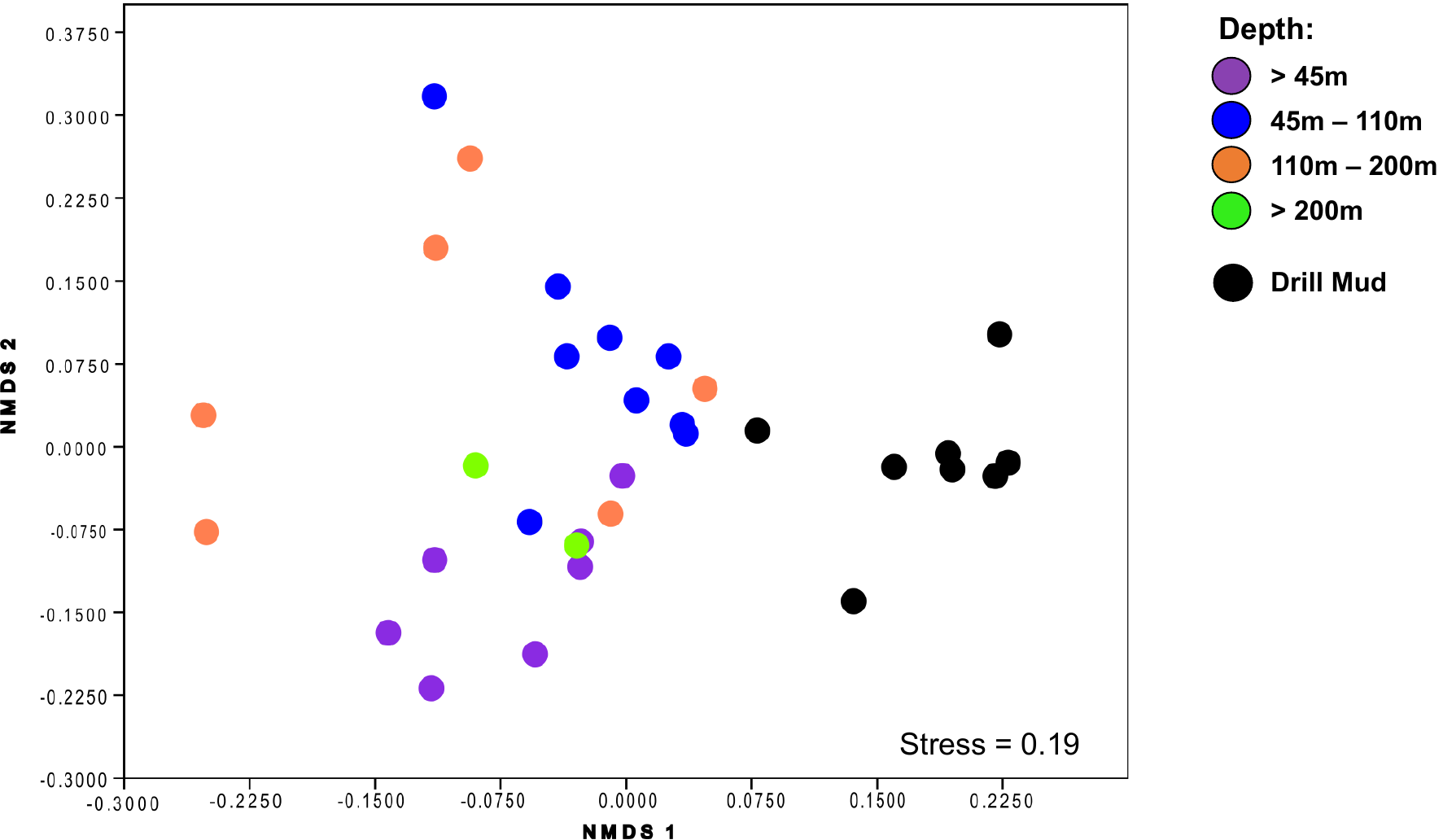

**Figure S2:** Nonmetric multidimensional scaling (NMDS) plot based on microbial community composition, depicting differences in microbial community structure across different drill core depths (color coded) and the analyzed drill core samples.

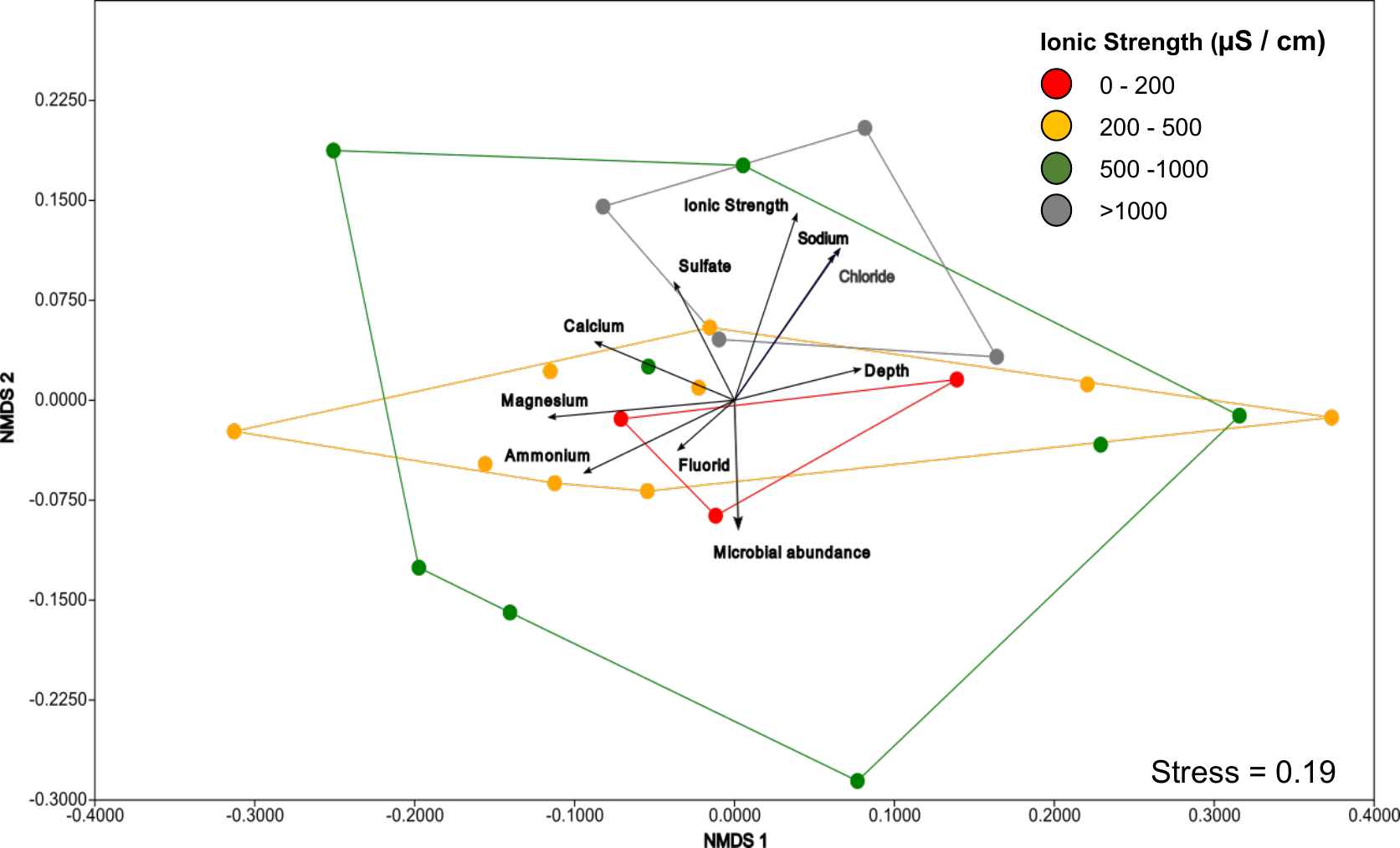

**Figure S3:** Nonmetric multidimensional scaling (NMDS) plot based on microbial community composition, depicting differences in microbial community structure across different ionic concentrations (color coded).

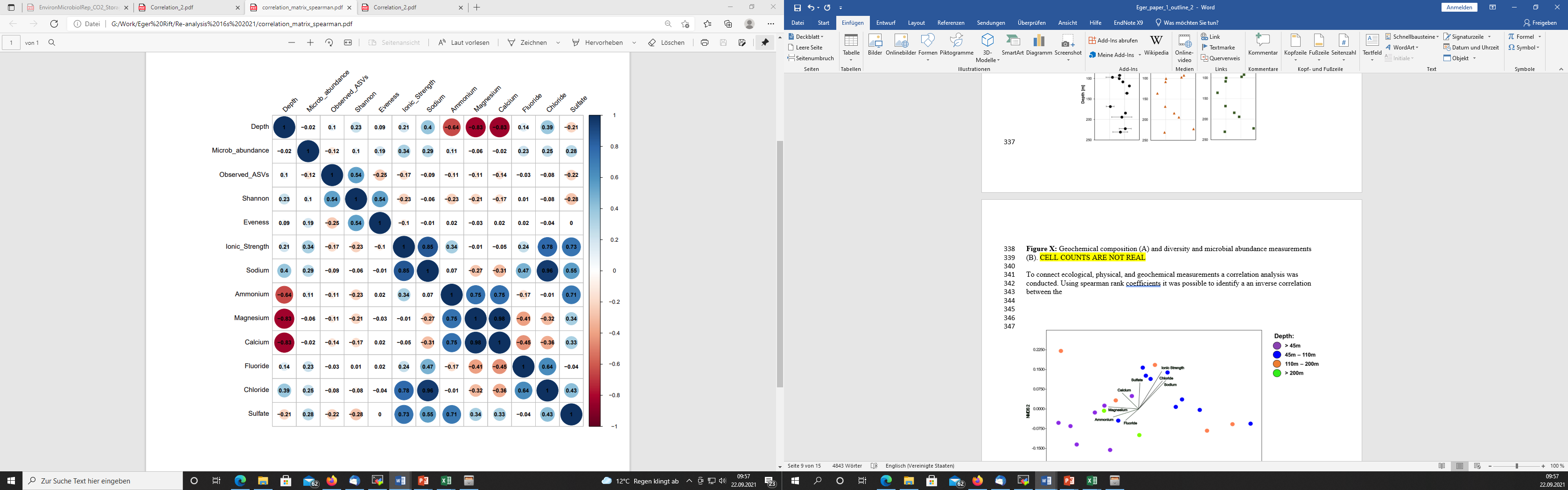

**Figure S4:** Heatmap showing spearman correlation coefficients between microbiological and geochemical measurements.

**Figure S5:** Distribution of the 25 most abundant microbial genera identified across the recovered drill core samples, color coded by class.

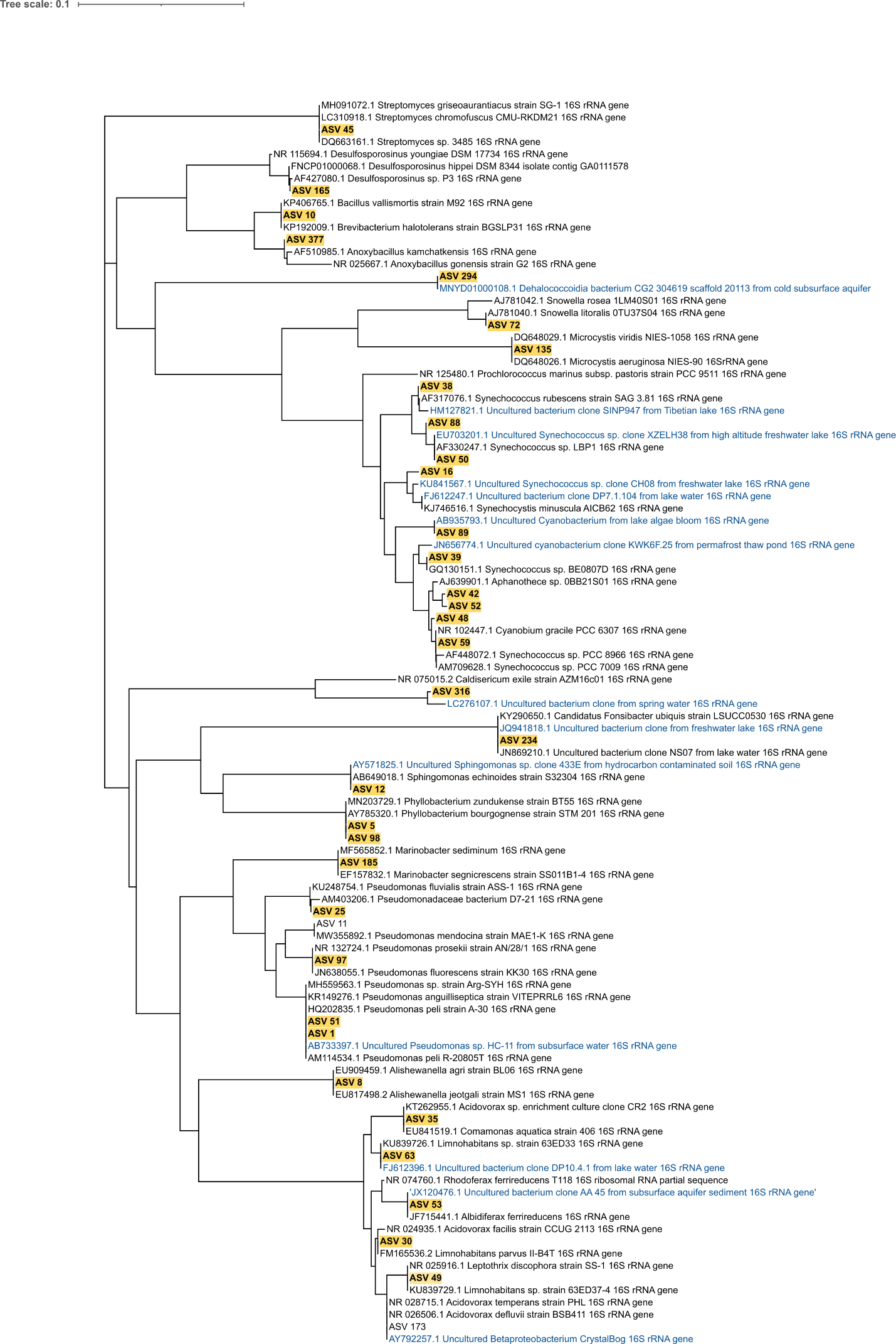

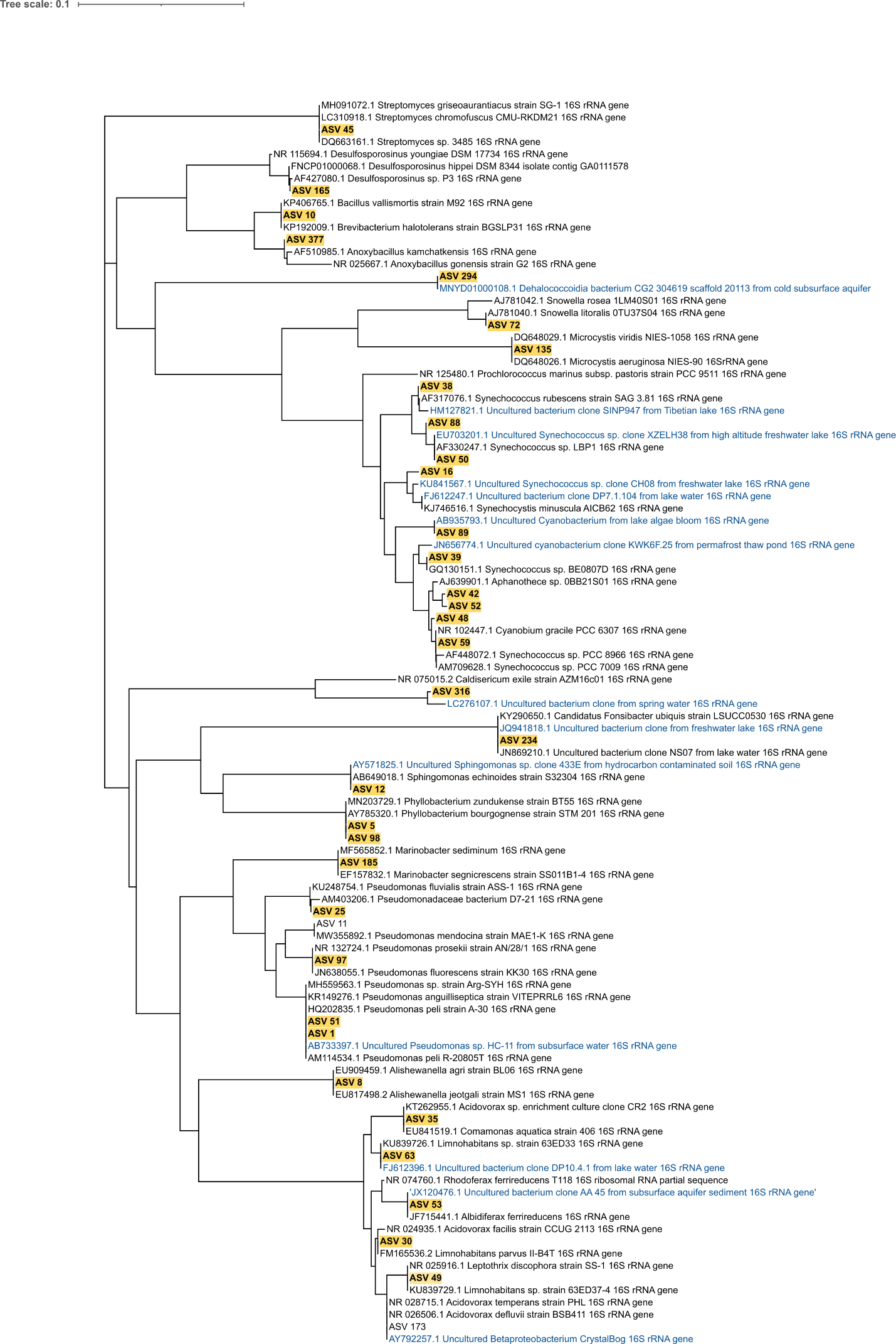

**Figure S6:** 16S rRNA Neighbor joining tree depicting the relationship between the Top 30 identified bacterial ASVs and their closest cultivated neighbors (black) and uncultivated sequences (blue)

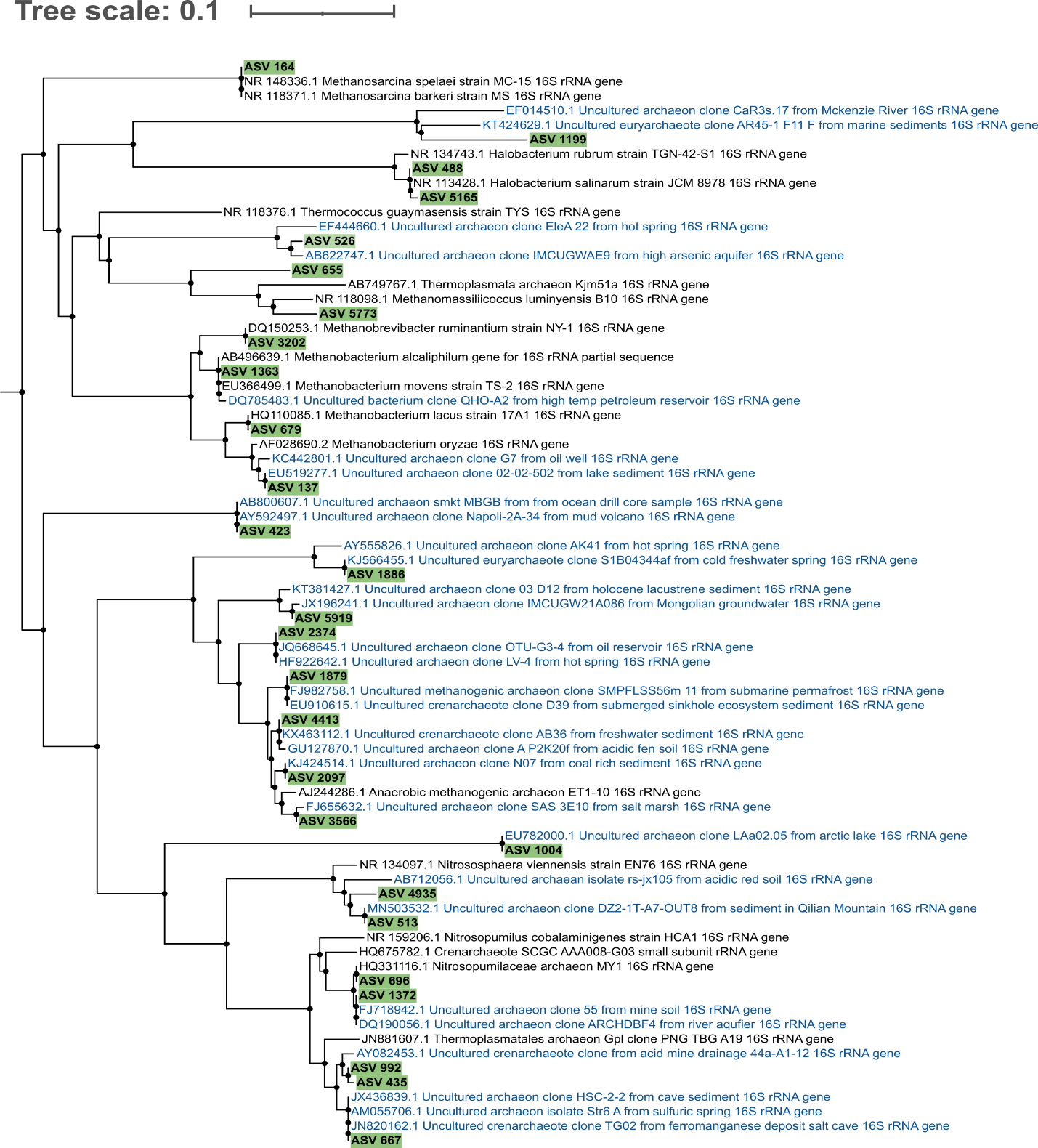

**Figure S6:** 16S rRNA Neighbor joining tree depicting the relationship between the Top 30 identified archaeal ASVs and their closest cultivated neighbors (black) and uncultivated sequences (blue)
